# Loss of nephronophthisis-associated nephrocystin-1 impairs DNA damage repair in kidney organoids

**DOI:** 10.1101/2025.06.09.658557

**Authors:** E. Sendino Garví, S. Biermans, N.V.A.M. Knoers, A.M. van Eerde, R. Masereeuw, M.J. Janssen, G.G. Slaats, A.M. van Genderen

**Author notes:** These authors contributed equally (M.J. Janssen, G.G. Slaats, A.M. van Genderen).

## Abstract

Nephronophthisis (NPH) is a heterogeneous, autosomal recessive ciliopathy and an important cause of end-stage renal disease (ESRD) in children and young adults. Since its classification as ciliopathy in 2003, NPH disease causal attribution had been focused primarily on ciliary dysfunction. The finding that ciliopathy players are involved in the DNA damage response (DDR) signaling resulted in a paradigm shift in thinking on NPH disease aetiology. Mutations in *NPHP1* are the leading cause of NPH, but the underlying mechanisms that lead to the disease phenotype remain poorly understood. Here, nephrocystin-1 depleted kidney organoids were generated and characterized to address this knowledge gap. We used CRISPR/Cas9 to generate *NPHP1* control (*NPHP1^WT^)* and two mutant (*NPHP1^ko1^* and *NPHP1^ko2^*) cell lines from healthy human induced pluripotent stem cells (iPSC), which then were differentiated into kidney organoids in an air-liquid interface following an optimized protocol. Upon loss of nephrocystin-1, kidney organoids showed impaired nephron structures and loss of glomerular mesangial and distal tubular cells. Furthermore, *NPHP1* depleted organoids exhibited a persistent inability to repair DNA lesions and showed increased senescence and fibrosis characteristics. Dynamic subcellular localization of nephrocystin-1 in *NPHP1^WT^*, particularly its translocation to nuclei 15 min post-UVC light exposure, suggested its direct involvement in the DDR. In conclusion, a novel *NPHP1*-depleted kidney organoid model was established, providing a platform to comprehensively study DNA damage, senescence and fibrosis simultaneously upon nephrocystin-1 loss. This advanced model aids in the understanding of the pathophysiology of NPH and paves the way towards identifying novel druggable targets.

## 1. INTRODUCTION

Nephronophthisis (NPH) is an autosomal recessive disease constituting the most predominant monogenetic cause for end-stage kidney disease (ESKD) in the pediatric and adolescent population [1]. Clinical manifestations of NPH encompass polyuria preceding a gradual decline in kidney function, attributable to progressive kidney scarring [2, 3]. Histological alterations include tubulointerstitial fibrosis, thickening and disruption of the tubular basement membrane, progressive atrophy of tubular epithelial cells, and in some cases the emergence of corticomedullary cysts in distal tubules [1, 2, 4]. Treatment of NPH primarily involves management of symptoms and complications, and ultimately kidney replacement therapy in cases of ESKD [5].

Over twenty genes have been implicated in NPH, with a substantial proportion of cases (approx. 50%) attributed to mutations in *NPHP1* encompassing both homozygous and compound heterozygous mutations, but mostly (approx. 80%) biallelic full gene deletions [1, 6, 7]. *NPHP1* encodes nephrocystin-1, a core protein situated in the transition zone at the base of the primary cilium within kidney tubule cells. This microtubule-based protrusion at the apical membrane plays a role in signaling pathways integral to kidney development and homeostasis, including Hedgehog, Wnt, and TGFβ pathways [2–4, 8]. Nephrocystin-1 orchestrates epithelial cell polarity through regulation of adherens junctions via the actin cytoskeleton, actin-binding proteins, and modulating proteins. Its deficiency is postulated to perturb the actin cytoskeletal network, leading to cyst formation and disruptions in nephron epithelial cell polarity [3, 4, 9]. Additionally, nephrocystin-1 deficiency is associated with inflammation, evidenced by upregulation of the chemokine CCL2, potentially contributing to tubulointerstitial fibrosis [4, 10].

Despite the advancements in NPH research in the last decade, the underlying mechanisms that lead to the progressive disease development remain unknown. A study in 2012 was the first to report a role for NPH-associated proteins in DNA damage response (DDR) signaling [11]. Later, another study showed the link between the loss of CEP290, a pan-ciliopathy protein, and an increase in replication stress-induced DNA damage [12]. Furthermore, activation of DNA damage signaling in NPH patient urine-derived cells was revealed, strengthening the previous indications [13]. Recently, a genome wide CRISPR screen in human retinal pigment epithelial cells showed that *NPHP1* knockouts sensitize the cells against formaldehyde and UV light [14], both treatments can create inter-strand crosslinks and single-strand lesions that would need to be removed by the nucleotide excision repair (NER) system [15]. NER is a key process for the removal of photolesions that, if left uncorrected, promotes replication stress [16], a condition in which DNA replication is disrupted. This includes stalling of the transcription machinery resulting in replication forks halting, which, if not promptly repaired, may lead to fork breakage and subsequent genomic instability [17]. Altogether, this evidence led us to hypothesize that *NPHP1* might be involved in DDR signaling and/or repair in the kidney which could contribute to the mitigation of replication stress.

Advancements in human *in vitro* models, particularly induced pluripotent stem cell (iPSC) technology coupled with CRISPR/Cas9 genome editing, offer a promising avenue for a more accurate advanced *in vitro* disease modeling. In this study, we aimed to generate and characterize CRISPR/Cas9-generated *NPHP1*-depleted kidney organoids utilizing human iPSCs. Furthermore, we studied the pathophysiological mechanisms underlying the disease phenotype upon *NPHP1* loss, with particular interest in DDR pathways.

## 2. RESULTS

### 2.1 Generation of kidney organoids depleted of nephrocystin-1

To investigate the impact of nephrocystin-1 depletion in kidney organoids, *NPHP1* knockout iPSC lines were obtained by targeting the *NPHP1* gene of a healthy iPSC parent line (*NPHP1^WT^*) using CRISPR/Cas9 technology (Figure 1A). After clonal expansion and sequencing, two iPSC clones containing *NPHP1* loss of function mutations (*NPHP1^ko1^* and *NPHP1^ko2^*) were selected for further analysis. Sanger Sequencing revealed *NPHP1^ko1^* to harbor a c.287_288insT mutation in one allele and a c.288_288delG mutation in the other allele (biallelic compound heterozygous) whereas *NPHP1^ko2^* contained the same c.287_288insT mutation in both *NPHP1* alleles (biallelic homozygous mutations) (Figure 1B). *In silico* predictions were used to estimate the effect of these mutations on the nephrocystin-1 protein based on a structure-based function prediction of the active site residues (Figure 1C). The nephrocystin-1^WT^ protein (eTM-score of 0.56+-0.15) was found to consist of 732 amino acids divided into five separate domains, of which domain 5 was predicted to contain the catalytic and protein-ligand binding residues (CscoreEC of 0.01 and 0.06, respectively) (Figure 1C). However, the premature stop codons generated by both mutations in the *NPHP1* genome, result in an amino acid sequence that only encodes part of domain 1, losing both the predicted catalytic and ligand binding residues of the protein when translated. As the termination codon of both mutations occurs >55 nucleotides upstream of the final intron/exon junction, the nonsense-mediated mRNA decay machinery is predicted to be activated leading to mRNA degradation before translation could occur. Therefore, both *NPHP1^ko1^*and *NPHP1^ko2^* are predicted to be full *NPHP1* knockouts without residual activity.

**Figure 1.**
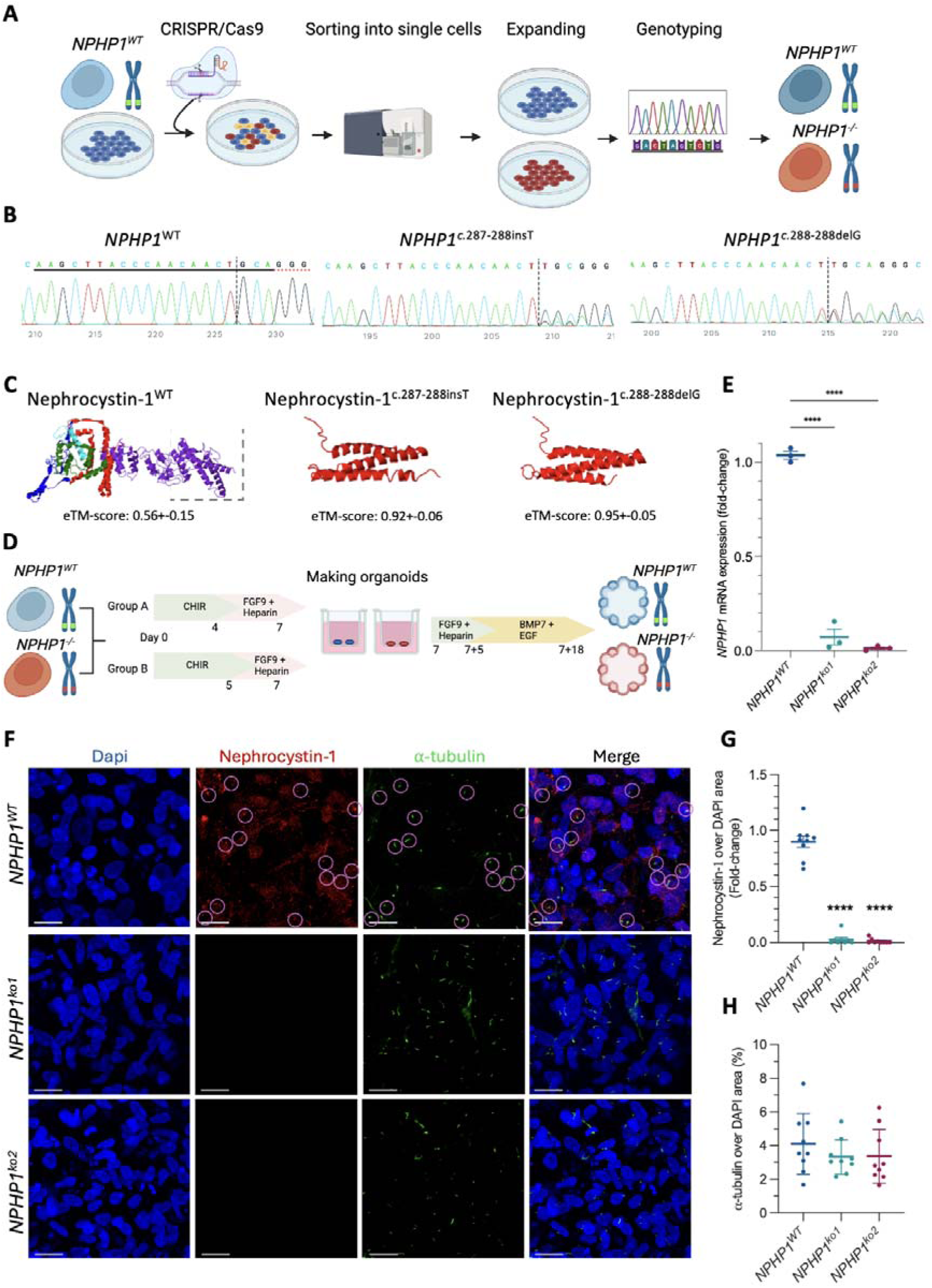
*NPHP1* depleted kidney organoids show loss of nephrocystin-1. **(A)** Schematic illustration of the disease model generation using iPSC and CRISPR/Cas9. **(B)** Sanger sequencing analysis of the target region confirmed that gene editing resulted in INDELs (insertion/deletions) of the *NPHP1* gene. The *NPHP1^-/-^*compound heterozygous line harbors a frameshift mutation in *NPHP1* (allele 1: c.287_288insT (predicted to introduce a premature stop codon: p.Gln101Ter), allele 2: c.288_288delG (predicted to introduce a premature stop codon: p.Leu112Ter)) and is referred to as *NPHP1^ko1^*. The *NPHP1^-/-^* homozygous line harbors a frameshift mutation in *NPHP1* (allele 1: c.287_288insT, allele 2: c.287_288insT (predicted to introduce a premature stop codon: p.Gln101Ter)) and is referred to as *NPHP1^ko2^.* **(C)** *In silico* prediction of the impact of premature stop codon on nephrocystin-1 protein sequence. The dashed square indicates the location of the active and binding domains in the WT nephrocystin-1. **(D)** Schematic illustration of the 25 days differentiation workflow from *NPHP1^WT^* and *NPHP1^-/-^* iPSCs to kidney organoids, including 7 days differentiation in 2D followed by 18 days maturation in liquid-air interface. (**E**) Quantification of *NPHP1* gene expression by qPCR of *NPHP1^ko1^* and *NPHP1^ko2^* compared to *NPHP1^WT^.* **(F)** Representative confocal images of the immunostainings against nephrocystin-1 (in red) and -tubulin (in green) in kidney organoids (Scale bar = 20 µm. Zoom in = 5X from the original image). The numerous pink circles indicate the presence of nephrocystin-1 at the base of the primary cilia in the *NPHP1^WT^*. **(G-H)** Image quantification of nephrocystin-1 and -tubulin in *NPHP1^WT^*, *NPHP1^ko1^* and *NPHP1^ko2^* organoids. One way ANOVA was performed (N=3, including 3 technical replicates for each biological replicate).

Kidney organoids were generated from both *NPHP1*-mutant iPSC lines and the WT line, and the presence and location of nephrocystin-1 was assessed with immunostainings (Figure 1D-H). To assess the predicted nonsense mediated decay, we measured the mRNA gene expression of *NPHP1*, confirming >95% depletion (Figure 1E). In WT conditions, nephrocystin-1 was detected in the cytoplasm, in the nucleus and at the basal body of the primary cilia (Figure 1F). Additionally, we quantified the depletion of nephrocystin-1 in both *NPHP1*-mutant organoids, observing a significant decrease of the protein expression levels of nephrocystin-1 of 95% and 99% in the *NPHP1^ko^*^1^ and the *NPHP1^k^*^o2^ kidney organoids, respectively (Figure 1F-G). We counterstained with -tubulin (Figure 1H) as nephrocystin-1 has been found co-localized in tubulin-rich structures including microtubules and centrosomes [18, 19]. Besides the primary cilia, but we found no clear co-localization nor differences in microtubules and/or primary cilia protein expression (Figure 1F&H).

### 2.2 Nephrocystin-1 depleted organoids show impaired nephron-like structures

To refine the potential role of nephrocystin-1 in the pathophysiology of NPH we assessed the presence of main nephron subpopulations in the organoids (Figure 2). We first evaluated the nephron structures presence, morphology and volumes of proximal tubules, distal tubules, glomeruli (including podocytes and mesangial cells) and distal segment of the distal tubule/collecting ducts (Figure 2). The nephron segment markers revealed that the *NPHP1^WT^* organoids exhibited contiguous nephron epithelia and segmentation into all 4 nephron segments, whereas *NPHP1^ko^*^1^ organoids showed a completely impaired physiological arrangement with loss of continuity between segments (Figure 2A). The *NPHP1^ko^*^1^ organoids failed to form tubular structures, resulting in bud-shaped proximal and distal tubules with empty spaces between all segments (Figure 2A). The *NPHP1^ko^*^2^ kidney organoid phenotype is presented as a multicellular aggregate consisting of predominantly distal segments and bud-shaped proximal tubules, rather than a discernable nephron-like phenotype (Figure 2A). To complement these findings quantitatively, volume quantification and transcriptomic analysis of *NPHP1^WT^*, *NPHP1^ko^*^1^ and *NPHP1^ko^*^2^ organoids were performed (Figure 1B). Overall, the volume of the different cell populations showed a trend towards a decrease in volume of proximal tubule and distal segments in *NPHP1^ko^*^1^ and a decrease in distal segments in *NPHP1^ko^*^2^ (Figure 2B). Complementary, our transcriptomic data confirmed an overall downregulation of the distal segment markers in both mutant models compared to *NPHP1^WT^*(Figure 2C). However, the expression of the distal segments, particularly the collecting duct gene expression marker, was already very low in *NPHP1^WT^* organoids (CPM<1) due to the differentiation protocol favoring towards more proximal segments (Table S1). To better assess organoid maturity, we further examined nephron progenitor and differentiation markers (*SIX2, PAX2, LIM1* and *SOX2*) (Figure 2C). In *NPHP1^ko^*^1^ organoids, *SOX2*, *PAX2* and *LIM1* were downregulated alongside *SIX2*, suggesting reduced nephron progenitor activity and impaired maturation. In contrast, *NPHP1^ko^*^2^ organoids showed upregulation of *SOX2*, *PAX2*, and *LIM1* with downregulation of *SIX2*. Altogether these results suggest that despite we can detect the presence of all nephron segments in all models, *NPHP1*-depleted organoids show less mature, poorly formed and connected structures, suggesting disturbed differentiation dynamics.

**Figure 2.**
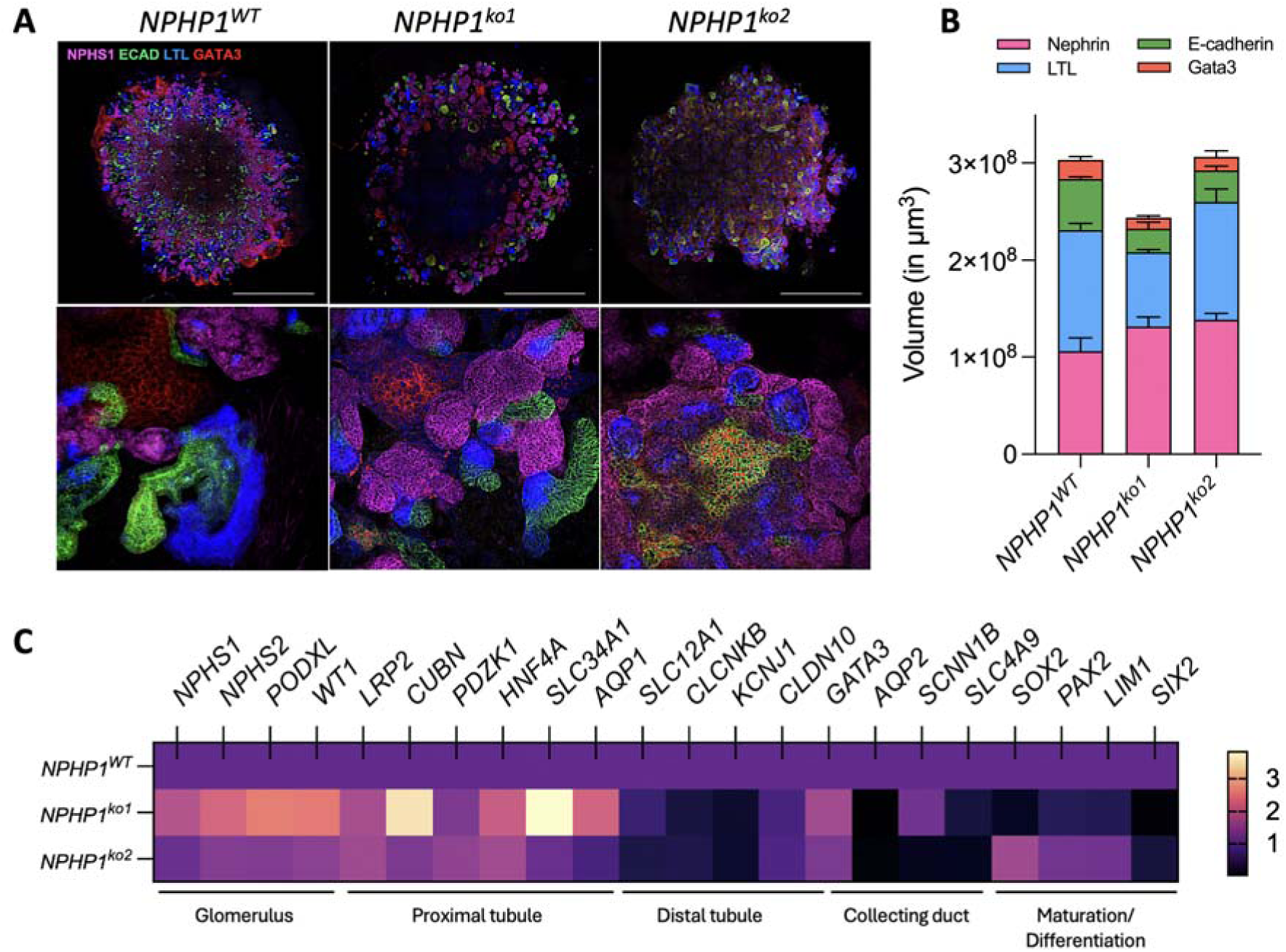
Mutant *NPHP1* kidney organoids show impaired nephron-like structures. (A) Representative tile scan immunofluorescence of kidney organoids displays impaired structural complexities between the *NPHP1^kos^*and the *NPHP1^WT^* kidney organoids. Each segment of the nephron is marked as: distal segment of the distal tubule/collecting duct (GATA3 + ECAD; red + green); distal tubule (ECAD; green); proximal tubule (LTL; blue); and glomeruli: including podocytes (NPHS1; magenta), and mesangial cells (GATA3; red). Scale bar = 1 mm. Zoom-ins on the bottom row = 10X from the top row. (B) Graphical presentation of the total volume (in µm^3^) of mutant and WT kidney organoids and the nephron segments immunostainings within each organoid. (C) Heatmap representation of fold-changes in count per million (CPM) of several markers for the different nephron segments in *NPHP1* mutant kidney organoids when compared to *NPHP1^WT^*, measured by RNA-sequencing. For panels B and C, each biological replicate represents measurement data from a single organoid of an independent differentiation round. N=3.

### 2.3 Nephrocystin-1 depleted organoids show alterations in DNA damage pathways

After filtering out transcripts with less than 20 counts per million (CPM) in our RNA sequencing data, PCA analysis revealed that upon loss of nephrocystin-1, both *NPHP1^ko^*^1^ and *NPHP1^ko^*^2^ organoids showed to be significantly different from the *NPHP1^WT^* (Figure 3A, Table S2). *NPHP1^ko^*^1^ and *NPHP1^ko^*^2^ organoids showed 635 and 488 downregulated genes; and 600 and 230 upregulated genes when compared to the *NPHP1^WT^*, respectively (Figure 3B, Table S3). Additionally, we compared the *NPHP1*-depleted organoids with each other and found 590 and 783 down- and upregulated genes, respectively (Figure 3B, Table S3). Enrichment upon differential expression analysis using KEGG revealed alterations in pathways related to DNA damage in *NPHP1^ko^*^1^ organoids when compared to *NPHP1^WT^*, including a 7-fold downregulation in DNA replication, a 4-fold downregulation in homologous recombination, in the Fanconi anemia pathway and in cell cycle regulation (Figure 3C, Table S4). Similarly, DNA-related pathways were identified when comparing *NPHP1^ko^*^2^ to *NPHP1^WT^*, including a ∼ 4-fold upregulation in DNA replication, and a 2-3-fold upregulation cell cycle, homologous recombination, and fanconi anemia pathways (Figure 3D, Table S5). A comparative analysis was performed between *NPHP1^ko^*^1^ and *NPHP1^ko^*^2^ to investigate the trancriptomic discrepancies between the two mutant organoid lines and our results showed that the top DEGs and enriched pathways were not related to any relevant pathway, particularly not related to DNA damage pathways (Figure S1, Tables S8-S11).

**Figure 3.**
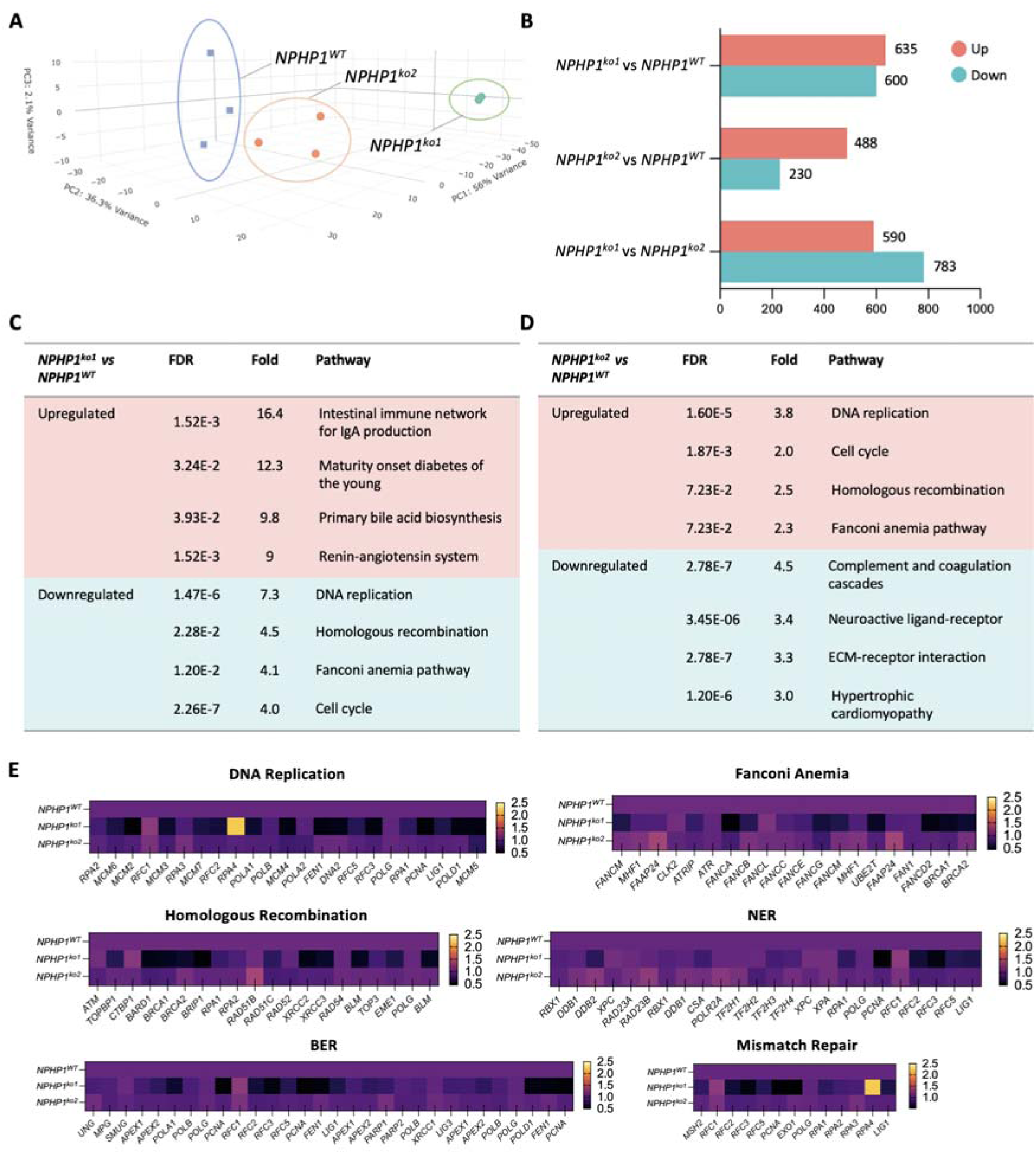
Transcriptomic analysis reveals alterations in DNA damage signaling pathways upon Nephrocystin-1 loss. (A) Principal component analysis (PCA) results represented in a 3-dimensional plot. (B) Quantification of the upregulated and downregulated genes after differential expression analysis. (C) Table presenting the top 4 most upregulated and most downregulated pathways after enrichment analysis when comparing *NPHP1^ko^*^1^ to *NPHP1^WT^*, shown in descending order based on their fold-change. (D) Table depicting upregulated and downregulated pathways after enrichment analysis when comparing *NPHP1^ko^*^2^ to *NPHP1^WT^*, shown in descending order based on their fold-change. (E) Heatmap presentation of fold-changes in count per million (CPM) of key genes in DDR pathways including DNA replication, Fanconi anemia, homologous recombination, nucleutide excision repair (NER), base-excition repair (BER) and mismatch repair pathways. The values plotted represent the fold-changes in *NPHP1* mutant kidney organoids compared to *NPHP1^WT^*. N=3 (except for *NPHP1^ko^*^1^, for which N=2). Each biological replicate represents a single kidney organoid of an individual differentiation round.

The observation of altered DDR signaling pathways in our *NPHP1^ko^*organoids is consistent with literature on the NPH-associated gene defects leading to impaired DDR [12, 20–22]. Next, we compared the expression changes amongst *NPHP1^WT^*, *NPHP1^ko^*^1^ and *NPHP1^ko^*^2^ organoids in several DDR pathways including DNA replication, Fanconi anemia, homologous recombination, nucleotide excision repair, base excision repair and mismatch repair (Figure 3E). Loss of nephrocystin-1 resulted in the overall downregulation of all DDR-related genes in *NPHP1^ko^*^1^ when compared to *NPHP1^WT^*, while *NPHP1^ko^*^2^ showed minor changes or an upregulation of all DDR-related genes (Figure 3E), confirming the results of the enrichment and pathway analysis in panels C and D. These findings indicate that without any stimuli, loss of nephrocystin-1 result in changes in DNA damage repair pathways.

### 2.4 *NPHP1* depleted organoids display differential BER and NER activity upon UVC light-induced DNA damage

To investigate whether nephrocystin-1 is involved in BER and/or NER activity upong DNA damage, we compared the expression changes amongst *NPHP1^WT^*, *NPHP1^ko^*^1^ and *NPHP1^ko^*^2^ organoids before and after the exposure to 25 J/m^2^ of UVC light (Figure 4). PCA analysis revealed strong clustering based on *NPHP1* genotype (Figure 4A, Table S6). All experimental conditions were then clustered (unsupervised clustering) based on the top 1000 differentially expressed genes (Figure 4B, Table S7), which confirmed that the clustering is mostly due to the *NPHP1* loss rather than UVC treatment.

**Figure 4.**
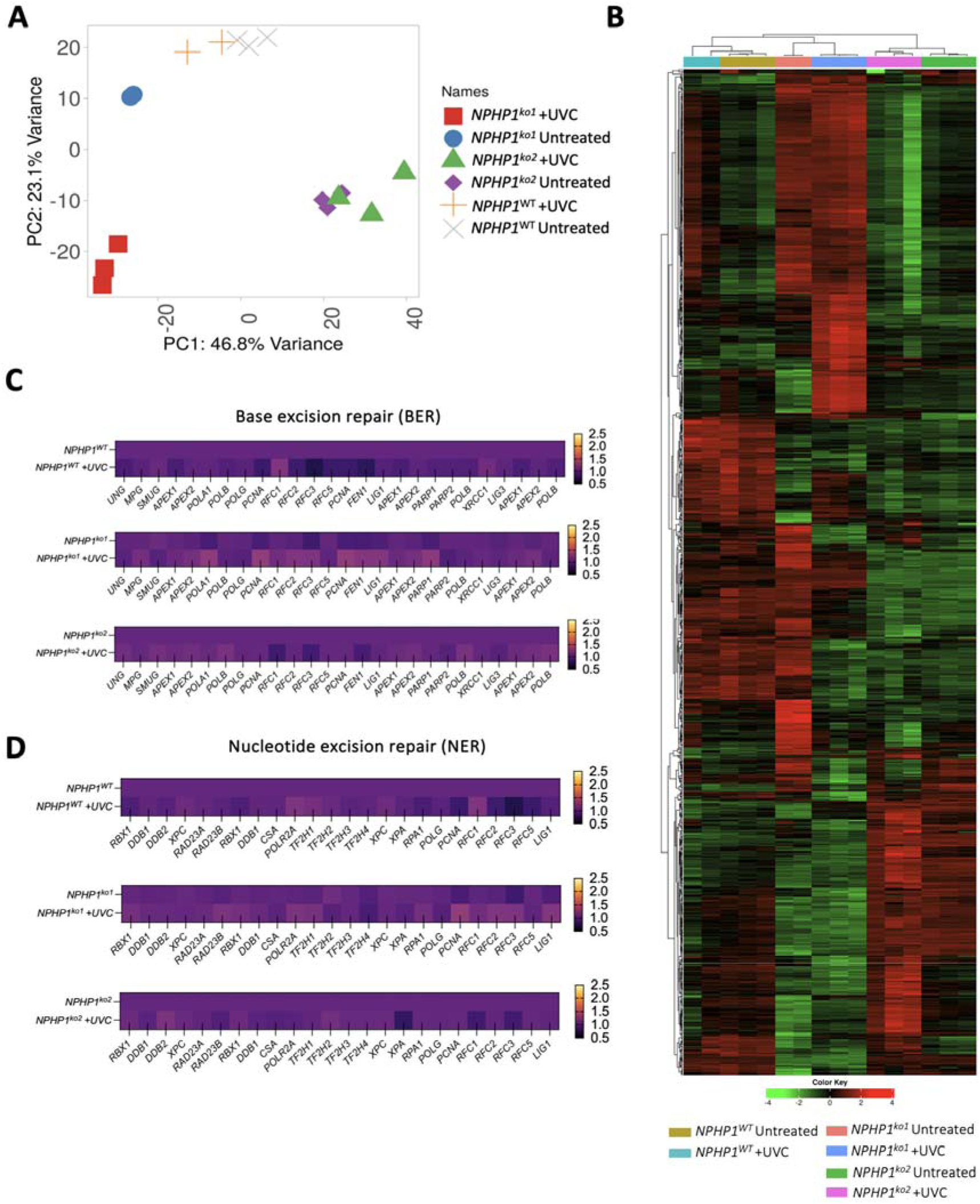
Transcriptomic analysis reveals alterations in base excision repair (BER) and nucleotide excision repair (NER) an DNA damage pathways upon nephrocystin-1 depletion after UVC exposure. **(A)** Principal component analysis (PCA) results. **(B)** Hierarchical clustering based on the top 1000 most differentially expressed genes after clustering for the samples. **(C-D)** Heatmaps presenting the fold-change of the CPMs of genes within the base excision repair (BER) pathway **(C)** and nucleotide excision repair (NER) pathways (**D)** of *NPHP1^WT^, NPHP1^ko^*^1^ and *NPHP1^ko^*^2^ exposed to UVC light, when compared to untreated conditions. All organoid lines were compared amongst each other under untreated conditions as well as their transcriptomic response to 15 min of 25J/m2 UVC light treatment. N=3, (except for *NPHP1^ko^*^1^ Untreated and *NPHP1^WT^ +* UVC, for which N=2), in which every biological replicate represents one single organoid.

To investigate the potential involvement of nephrocystin-1 in DDR under healthy conditions, as well as the effect of the UVC treatment on these pathways upon nephrocystin-1 loss, we focused our transcriptomic analysis on the gene expression patterns of key genes within the NER and BER pathways pathways (Figure 4C-D). After 15 min of exposure to 25 J/m^2^ of UVC light, *NPHP1^WT^* organoids engage both NER and BER, upregulating and downregulating different subsets of genes within both pathways. In contrast, *NPHP1^ko^*^1^ organoids mostly upregulate NER and BER while the same genes show either minor up and down regulation or remained unchanged in *NPHP1^ko^*^2^ organoids. These results suggest that nephrocystin-1 loss might impact the correct functioning of NER and BER pathways.

### 2.5 Nephrocystin-1 accumulates in the nucleus upon UVC light exposure and is essential for DNA lesion repair

Exposure to UVC light induces two types of photolesions: 6-4 photoproducts (64PPs) and cyclobutane pyrimidine dimers (CPDs), both with a high mutagenic potential. To confirm the role of nephrocystin-1 in DNA damage repair (specifically NER and BER pathways, as our results suggested earlier), we investigated the clearance of 64PPs and CPDs after UVC light exposure in addition to the expression and localization of nephrocystin-1 in cells within the *NPHP1^WT^* kidney organoids (Figure 5). Prior to UVC light exposure, we did not detect the presence of 64PPs or CPDs in any of the lines, except for an insignificant presence of 64PP in the unexposed *NPHP1^ko^*^1^ (Figure 5 A-D). After 15 min of UVC light exposure, *NPHP1^WT^* kidney organoids showed an increase in both 64PPs and CPDs. Both *NPHP1^ko^*^1^ and *NPHP1^ko^*^2^ accumulated 64PPs and CPDs, with lower amounts of 64PPs in the *NPHP1^WT^*, while the presence of CPDs was similar in the mutant models. The *NPHP1^WT^* was able to clear 64PPs lesions within the first 2 h after exposure while both *NPHP1^ko^*^1^ and *NPHP1^ko^*^2^ presented 64PPs even after 24 h (Figure 5C). Similarly, *NPHP1^WT^* organoids cleared all their CPDs lesions within the first 8 h after exposure while *NPHP1^ko^*^1^ and *NPHP1^ko^*^2^ still had some remaining CDPs after 8 up to 24 h (Figure 5D). At 15 minutes post-UVC exposure, both *NPHP1*^ko1^ and *NPHP1*^ko2^ organoids showed reduced detection of 64PPs and CPDs compared to *NPHP1^WT^*. Since photolesions form instantaneously upon UVC exposure, this reduction is unlikely to reflect true differences in lesion formation. Instead, this can be interpreted as reduced lesion detectability due to impaired chromatin remodeling in nephrocystin-1 deficient cells, which may delay antibody accessibility to damage sites. At later timepoints, photolesion signals accumulate and persist longer in *NPHP1*^ko1^ and *NPHP1*^ko2^ than in WT, consistent with delayed recognition and repair initiation. This suggests that nephrocystin-1 might play a role in facilitating chromatin relaxation.

**Figure 5.**
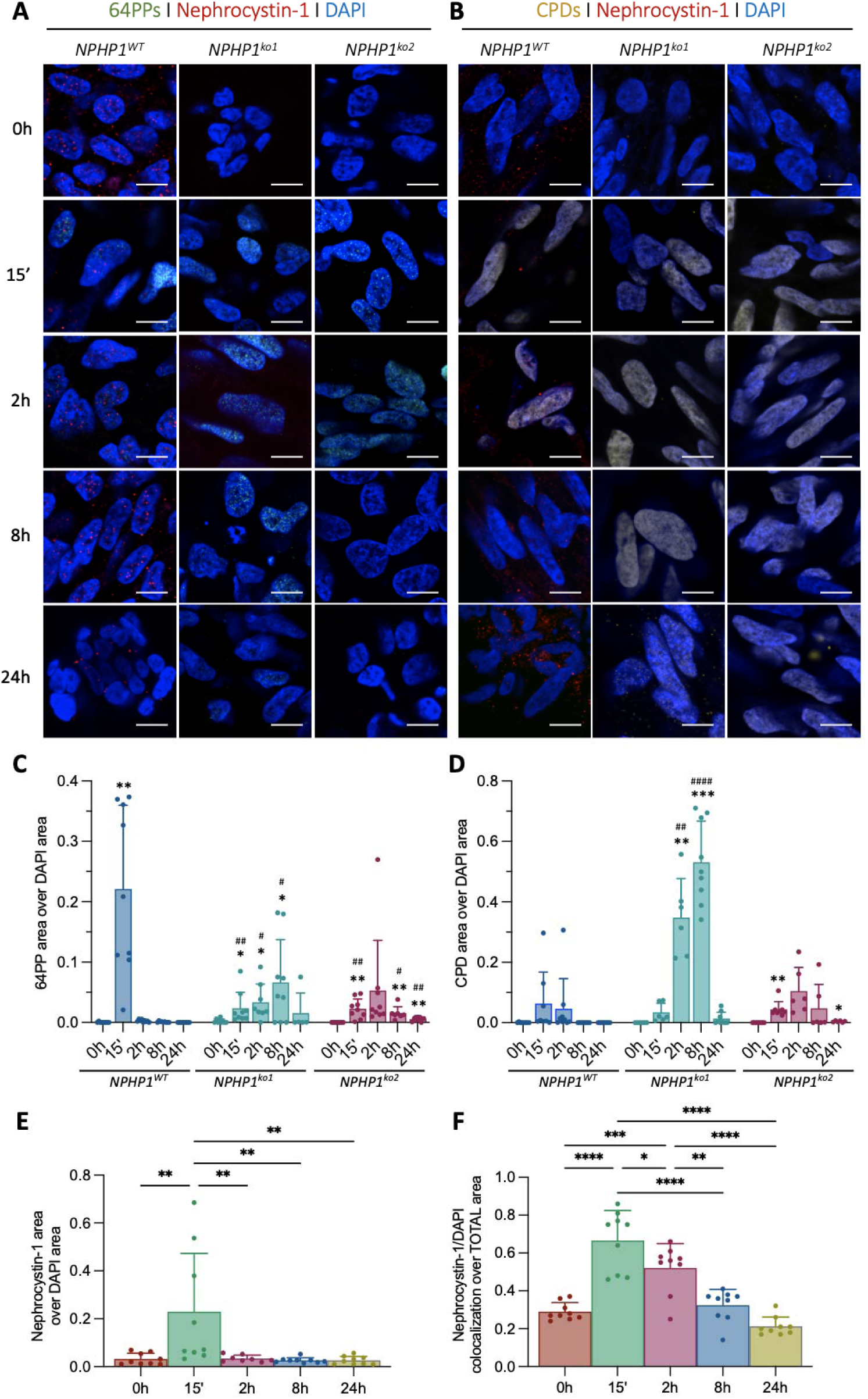
The ability to repair 64PP and CPDs upon UVC light exposure is decreased upon nephrocystin-1 loss. **(A-B)** Representative immunostainings at 63X plus 5X digital zoom of 64PPs (**A)** and CPDs (**B)**, in addition to nephrocystin-1 in all three organoid lines at different timepoints after exposure to 25 J/m^2^ of UVC light. Unmerged channels for each datapoint in panels A and B can be found in Figures S2 and S3, respectively. (**C-D)** Image quantification of the 64PPs and CDPs total area against the total DAPI area. (**E)** Image quantification of the total nephrocystin-1 area in *NPHP1*^WT^ organoids within the image normalized by DAPI area. (**F)** Quantification of the nephrocystin-1 area within the nuclei localization (DAPI), normalized by the total nephrocystin-1 area in the image. For C and D, a two-way ANOVA analysis with multiple comparisons analysis was performed (N=3, including 3 technical replicates for each biological replicate. * represent significance when comparing timepoints within the same cell line and # represent significance when comparing the same timepoint amongst the different cell lines). For E and F, a one-way ANOVA analysis was performed (N=3, including 3 technical replicates for each biological replicate). Scale bar = 10 µm.

Furthermore, we used the *NPHP1^WT^* kidney organoids to investigate the location and behavior pattern of nephrocystin-1 at various timepoints after UVC light exposure to assess whether the protein is involved in response to DNA lesions (Figure 5E-F). Our results show accumulation of nephrocystin-1 15 min after UVC light exposure, which returned to baseline levels within 2 h. Additionally, when we measured the presence of nephrocystin-1 solely in the nuclear fraction, we found an increase in the nuclei 15 min after UVC light exposure, which gradually restored over the following 24 h until reaching resting conditions. Overall, these results show that within the first 15 min after UVC light exposure, 64PPs and CPDs accumulated in the nuclei of *NPHP1^WT^* while the presence of nephrocystin-1 increased, particularly in the nuclei. Furthermore, 64PPs and CPDs were still detectable in both *NPHP1^ko^*^1^ and *NPHP1^ko^*^2^, while they were all cleared within the first 2 h in the *NPHP1^WT^*. These findings suggest that nephrocystin-1 plays a role in the DDR upon small DNA lesions.

### 2.6 *NPHP1* depleted kidney organoids show increased DNA damage signaling

Several of the NPH-associated proteins have been shown to be involved in the DDR signaling and replication stress response [11, 12]. The phosphorylation of histone 2AX (γH2AX) occurs in response to DNA damage, and it is an early indicator of DNA double strand breaks [23]. Although UVC exposure does not directly induce dsDNA breaks, the failure to clear 64PP and CPDs in a timely manner will lead to dsDNA breaks and consequent replication fork arrest. Therefore, we evaluated the presence and clearance of DNA damage using γH2AX as a marker (Figure 6) and we quantified γH2AX at later timepoints after UVC light exposure to evaluate dsDNA break clearance. Our results showed the presence of γH2AX in both *NPHP1^ko^*^1^ and *NPHP1^ko^*^2^ prior to UVC light exposure, indicating accumulating dsDNA damage under untreated conditions (Figure 6A-B). The *NPHP1^WT^* accumulated γH2AX for the first 8 h which returned to baseline levels after 24 h, while the *NPHP1^ko^*^1^ and *NPHP1^ko^*^2^ were unable to remove γH2AX even 3 days after exposure (Figure 6A-B).

**Figure 6.**
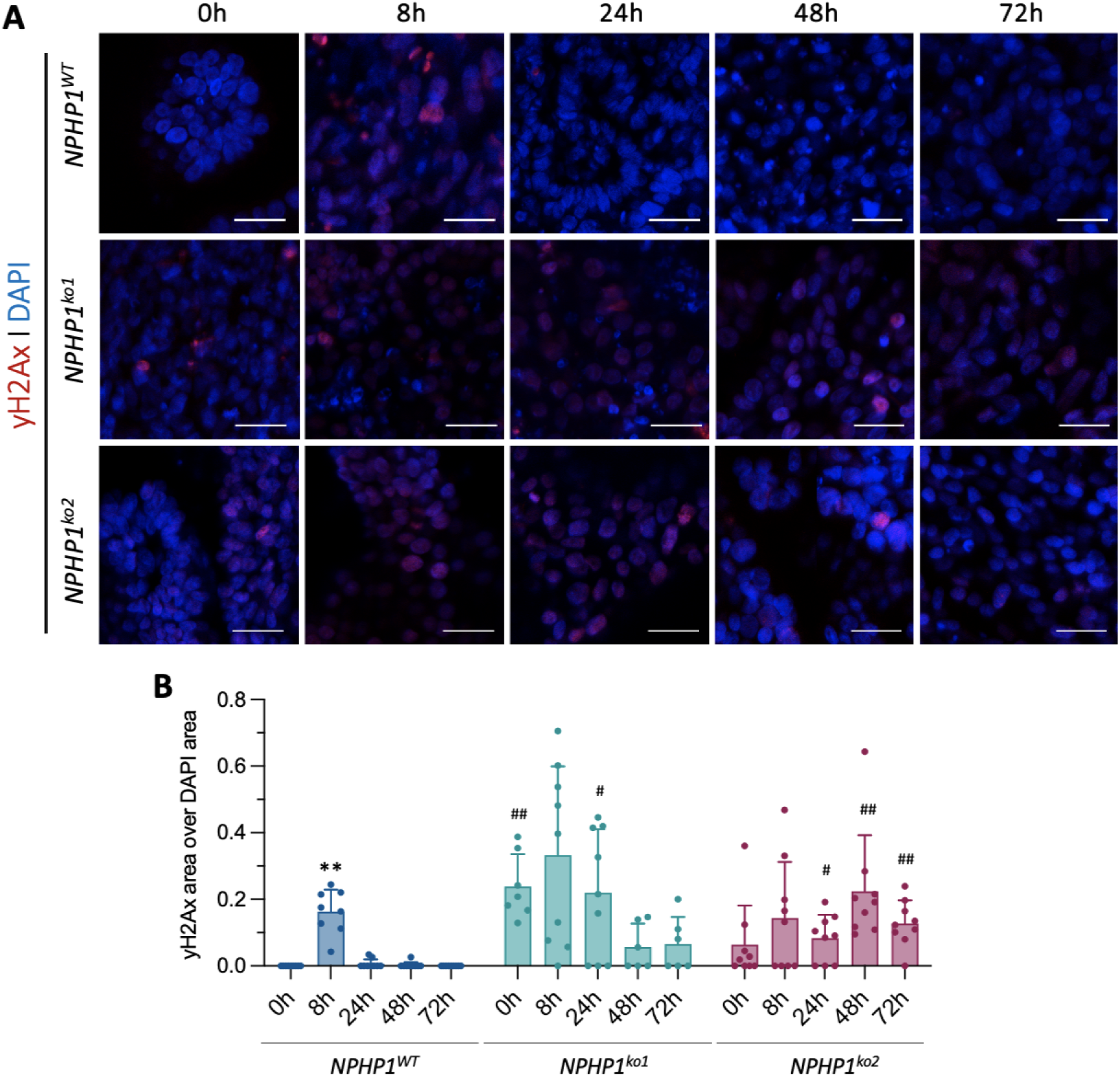
dsDNA damage signalling accumulates in *NPHP1* depleted organoids. **(A)** Representative immunostaining images at 63X plus 3X digital zoom of γH2AX in all three organoid lines at different timepoint after exposure to 25 J/m^2^ of UVC light. (**B)** Quantification analysis of the γH2AX area over DAPI area of the panels above. A two-way ANOVA analysis with multiple comparisons analysis was performed (N=3, including 3 technical replicates each. The * represent significance when comparing timepoints within the same cell line and the # represent significance when comparing the same timepoint amongst the different cell lines). Scale bar = 20 µm.

To ameliorate the observed γH2AX accumulation upon nephrocystin-1 loss, we hypothesized that arresting the cell cycle at the G2/M boundary would allow the cells to repair the DNA damage we observed in the *NPHP1^ko^*^1^ and *NPHP1^ko^*^2^. Therefore, we treated all organoids with 10 nM of RO-3306, a potent selective inhibitor of CDK1 that has recently been described to prevent mitotic entry and arrest cell division at the G2/M boundary [24] (Figure S4). Our results show that despite RO-3306 treatment did not improve yH2Ax presence in *NPHP1^ko^*^2^, it was able to reduce the γH2AX accumulation in the *NPHP1^ko^*^1^ after 24 h of treatment (Figure S4).

### 2.7 Kidney organoids show accumulation of senescent cells and pro-fibrotic markers upon nephrocystin-1 loss

The continuous presence of DNA damage will activate the DDR pathways for which a sustained activation promotes cell-cycle arrest and induces an irreversible senescent state in kidney cells [25, 26]. One of the phenotypes observed in NPH patients is the presence of senescent cells in the kidney [27]. To investigate whether the loss of nephrocystin-1 would lead to cellular senescence, we measured β-galactosidase activity in kidney organoids (Figure 7A-B). Immunostaining results show a significant increase in β-galactosidase positive cells in both the *NPHP1^ko^*^1^ and *NPHP1^ko^*^2^ kidney organoids, when compared to the *NPHP1^WT^*organoids (Figure 7A-B), indicating a widespread senescence state in the kidney organoids upon nephrocystin-1 loss.

**Figure 7.**
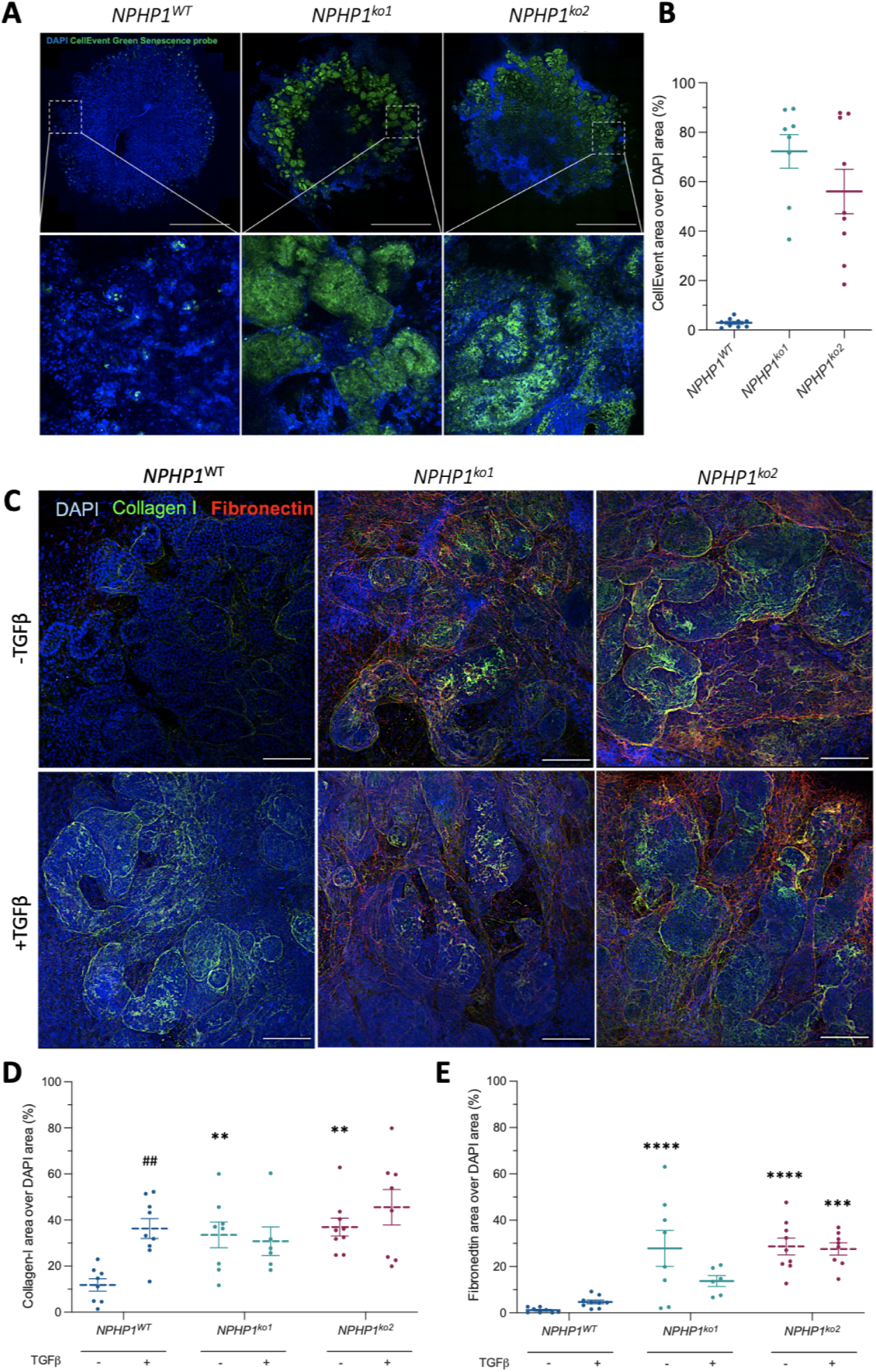
Senescent cells and fibrosis-associated proteins accumulate upon nephrocystin-1 depletion. **(A)** Representative immunostainings of tile scans of entire organoids using the CellEvent probe as marker for β-galactosidase activity. Scale bar = 1 mm. Zoom = 10X. **(B)** Quantification of the senescence immunofluorescent images shown as CellEvent area over total DAPI area. **(C)** Representative immunostainings of all three organoids using collagen-I and fibronectin as markers for fibrosis. **(D-E)** Quantification of the immunofluorescent images of collagen-I (**D)** and fibronectin (**E)** shown as area over total DAPI area. TGFβ was added at a final concentration of 50ng/mL for 24 h. For B, a one-way ANOVA analysis was performed (N=3, each representing 3 technical replicates, p <0.05). For D and E, a two-way ANOVA analysis with multiple comparisons analysis was performed (N=3 biological replicates, each representing 3 technical replicates, p <0.05). Scale bar = 100 µm.

Kidney senescence has been linked to fibrosis progression [28] therefore, we also assessed the presence of the extracellular matrix proteins collagen-I and fibronectin with and without pro-fibrotic stimulation with TGFβ (Figure 7C-E). Without any pro-fibrotic environment stimulation, both *NPHP1^ko^*^1^ and *NPHP1^ko^*^2^ kidney organoids showed a significant increase in protein expression of collagen-I and fibronectin when compared to the *NPHP1^WT^* organoids (Figure 7D-E). TGFβ stimulation induced a significant increase in collagen-I deposition in the *NPHP1^WT^* organoids but had no significant effect on either of the *NPHP1*-mutant organoids (Figure 7D), suggesting that the fibrotic response in the *NPHP1*-mutant organoids probably was already saturated, as observed in the untreated conditions. Pathway analysis (KEGG) on our RNA-Seq data showed that despite differential up and down regulation of key players, senescence and pro-fibrotic pathways were not significantly affected in the *NPHP1*-mutant organoids (Figures S5-8). However, we observed differential regulation of key players within the canonical TGFβ pathway in *NPHP1*^ko1^ and *NPHP1*^ko2^ organoids. In *NPHP1*^ko1^, upstream components of the TGFβ cascade were downregulated, while downstream SMAD-dependent targets were upregulated. Similarly, in *NPHP1*^ko2^ the upstream TGF-β signaling was downregulated while SMAD-dependent targets remained unchanged. Furthermore, non-canonical MAPK and PI3K/Akt pathways were engaged in both *NPHP1*^ko1^ and *NPHP1*^ko2^ (Figures S9-S12). Together, these findings suggest that loss of nephrocystin-1 leads to an altered responsiveness to TGFβ stimulation, with impaired canonical signaling and possible compensation via non-canonical pathways.

## 3 DISCUSSION

In this study, we modeled *NPHP1*-mediated NPH in human iPSC-derived kidney organoids. By applying CRISPR/Cas9 gene editing in iPSCs, we obtained *NPHP1* depleted kidney organoid models that accurately recapitulate multiple aspects of the NPH phenotype, including the impairment of nephrons, presence of senescent cells and fibrotic markers. Moreover, we used these *NPHP1* disease models to further study the pathomechanism of NPH and to elucidate the causal link between nephrocystin-1 loss and the disease phenotype. Our results strongly indicate that nephrocystin-1 is involved in the DDR.

Ciliary components can travel to several cellular compartments, including the nuclei, upon cilia motion sensing or during cell division [29–31]. We found nephrocystin-1 to be mostly located in the cytoplasm of the cells in the *NPHP1^WT^* in resting conditions, which translocated to the nuclei of the cells within 15 min after UVC light exposure, strongly indicating that nephrocystin-1 plays an extra-ciliary role in DDR, either in DNA damage recognition, DNA repair machinery recruitment (signaling), or DNA damage repair. Our transcriptomic results show that the cellular response to DNA damage and the DNA repair pathways were upregulated upon loss of nephrocystin-1. While *NPHP1^ko1^*and *NPHP1^ko2^* differ in the specific sets of dysregulated genes, they converge on the impairment of the same DNA damage response pathways. Response to DNA damage, including recognition and repair are very dynamic processes and there is a possibility that at transcriptomic level, we did not capture the true *NPHP1*^ko2^ response to DNA damage, as the transcriptomic profiles of *NPHP1*^ko1^ and *NPHP1*^ko2^ suggest non-identical pathway compensation. Experimental data confirmed an impaired response to DNA damage in *NPHP1*^ko1^ and *NPHP1*^ko2^, as well as increased yH2Ax signaling, which confirms DDR pathways impairment, although with less sensitivity than *NPHP1*^ko1^. While immunostaining for NPHP1 showed no detectable signal in either *NPHP1*^ko1^ and *NPHP1*^ko2^, western blot analysis revealed a faint residual band in *NPHP1*^ko2^ (Supplementary Figure S13), indicating that this clone may retain low-level expression of a NPHP1 protein product. This suggests that the observed divergenge reflects a combination of clonal adaptation and residual protein expression in *NPHP1*^ko2^, whereas the shared DDR impairment underscores the reproducibility and biological relevance of our *NPHP1* models.

While *NPHP1^WT^* organoids resolved 64PPs and CPDs derived from UVC light stress within 2 h post-exposure, both *NPHP1* depleted organoid models failed to eliminate these photoproducts even up to 24 h after exposure. We unexpectedly detected lower photolesion signal at the 15’ timepoint in *NPHP1^ko1^* and *NPHP1^ko2^*organoids compared to the *NPHP1^WT^*, which can be explained by delayed chromatin remodeling rather than reduced lesion formation. *NPHP1* loss may impair the rapid chromatin relaxation normally triggered by UVC damage, thereby reducing early antibody accessibility to lesions. This interpretation is supported by the subsequent persistence of photolesions in *NPHP1^ko1^* and *NPHP1^ko2^* organoids, highlighting a role for nephrocystin-1 in facilitating early DDR recognition events. Additionally, *NPHP1^ko1^* and *NPHP1^ko2^*organoids differ in the extent of CPD accumulation, despite both lacking nephrocystin-1. As shown in Figure 2, proximal tubule volumes (where loss of nephrocystin-1 should have a bigger impact) were present and comparable across all organoid lines, indicating that differentiation efficiency is unlikely to explain this variability. Instead, these differences most likely reflect distinct adaptations to the cloning process and the distinct CRISPR-induced mutations in the two knockout lines. The *NPHP1*^ko1^ line carries two different frameshift mutations, c.287_288insT (p.Gln101Ter) in allele 1 and c.288_288delG (p.Leu112Ter) in allele 2, whereas *NPHP1*^ko2^ carries the same c.287_288insT frameshift in both alleles. Because the lines share the c.287_288insT mutation but differ in the second allele of *NPHP1*^ko1^, it is possible that the c.288_288delG mutation results in a distinct adaptive state that increases susceptibility to DNA damage-related defects. Transcriptomic analyses (Figures 2C and Figure 3) support this interpretation, showing enrichment of DDR pathways in both *NPHP1*-depleted lines but with gene-level differences suggestive of alternative coping strategies. Importantly, both lines converge on shared DDR hallmarks, delayed lesion recognition, persistent photolesions, prolonged γH2AX signaling, and downstream senescence and fibrosis, underscoring the robustness of the model. Such clonal variation may even parallel the phenotypic heterogeneity observed among NPH patients [32].

Interestingly, the increased detection of nephrocystin-1 in the nuclei appeared to correlate with 64PP and CPDs detection and clearance, further supporting the role of nephrocystin-1 in DDR. Although nephrocystin-1 localizes to primary cilia and adherens junctions under basal conditions, our data does not support the idea that the nuclear pool originates from these structures. The observed nuclear enrichment after UVC exposure could also be consistent with an enhanced nuclear translocation from the cytoplasm from de novo protein synthesis, or from a combination of all known locations in the cell.

Upon loss of nephrocystin-1, kidney organoids stained positive for γH2AX with no additional treatment. The γH2AX signal observed in NPHP1-deficient organoids likely represents both focal DSBs [33] and pan-nuclear γH2AX associated with apoptosis or advanced senescence [34]. Pan-nuclear γH2AX has been reported as a hallmark of apoptosis, resulting from ATM/ATR-dependent hyperphosphorylation of H2AX across the entire nucleus during chromatin condensation and fragmentation. This interpretation is consistent with the accumulation of persistent lesions, DNA damage signaling, and senescence we observed in *NPHP1-*deficient organoids. It is also important to note that UVC light exposure generates dsDNA breaks (DSB) if the photolesions caused upon exposure are left uncorrected, which can consequently lead to replication stress [35–37]. Therefore, it could be hypothesized that, if nephrocystin-1 has a role in DDR, it will still be detectable in the nuclei of the cells in dsDNA damage. However, the detected nephrocystin-1 in the nuclei decreased between 2 - 8 h after UVC treatment while γH2AX was still detectable in the *NPHP1^WT^* organoids, suggesting that nephrocystin-1 might not have a direct effect on dsDNA damage repair but rather through signaling.

Upon loss of nephrocystin-1, the kidney organoids were positive for γH2AX, indicating the presence of dsDNA damage under untreated conditions. Both *NPHP1* depleted organoids were unable to reduce dsDNA damage and/or ameliorate replication stress caused by treatment with UVC light, in contrast to the repair observed in *NPHP1^WT^* within the first 2 h after UVC light stress. Additionally, we aimed to arrest cell cycle in the G2/M boundary to give the cells more time to repair the DSB and/or replication stress, which was achieved in the case of *NPHP1^ko1^* by a 24 h treatment with a well-known potent cell cycle inhibitor (RO-3306) without triggering apoptosis [24, 38]. However, *NPHP1^ko2^* did not respond as treatment with RO-3306 did not reduce yH2Ax detection. We observed that *NPHP1^ko2^* organoids exhibit lower basal γH2AX levels compared to *NPHP1^ko1^*, suggesting a distinct DDR or cell cycle state that may blunt responsiveness to G2/M arrest. Additionally, clonal variability in DNA damage recognition and repair kinetics, as discussed earlier, could also influence RO-3306 sensitivity. Lastly, if *NPHP1^ko2^*cells are already partially arrested in G2/M due to replication stress, RO-3306 would be less effective.

Despite the gap in the understanding of the exact molecular mechanism of nephrocystin-1 loss in NPH it is known that DNA damage increases replication stress, which could in turn put the cells into a senescent state, which eventually contributes to the kidney fibrotic phenotype found in patients [39–41]. The phenotypical characterization upon nephrocystin-1 loss confirmed increased levels of senescence in the tubular structures of our *NPHP1^-/-^* kidney organoids compared to the *NPHP1^WT^*. Interestingly, kidney fibrosis has been correlated with arrest of kidney tubular epithelial cells in the G2/M phase of the cell cycle, the universal marker of cellular senescence [42, 43]. Upon irreversible cellular arrest, which could be induced by a high DNA damage and subsequent replication stress, tubular epithelial cells can activate JNK signaling to induce pro-fibrotic factors and promoting fibroblast activation and fibrosis [42, 43]. Besides, continuous secretion of pro-fibrotic factors by the senescent cells causes EMT in kidney tissues, contributing to fibrosis and collagen deposition after reprogramming, entering a vicious cycle [42, 44]. Moreover, we also observed significantly higher levels of fibronectin and collagen-I deposition in our mutant organoids compared to their isogenic control. Collagen type I signal in *NPHP1* depleted kidney organoids localizes predominantly to the basal side of tubular structures, with enrichment consistent with pathological fibrosis, including accumulation near or within tubular basement membranes [45]. However, we observed no response to TGFβ stimulation in *NPHP1^-/-^* kidney organoids, while we did in *NPHP1^WT^*. TGFβ is a key fibrosis molecule that induces epithelial-to-mesenchymal transition (EMT), contributing to tubulointerstitial fibrosis [46, 47]. The absence of further induction of fibrotic markers by TGF-β stimulation in mutant organoids may reflect both saturation of ECM deposition and altered signaling responsiveness. Our transcriptomic analysis revealed genotype-specific dysregulation of the canonical TGFβ/SMAD pathway in *NPHP1*^ko1^ and *NPHP1*^ko2^, with aberrant downstream activation in *NPHP1*^ko1^ and reduced responsiveness in *NPHP1*^ko2^. Additionally, both *NPHP1*^ko1^ and *NPHP1*^ko2^ exhibited the engagement of non-canonical MAPK and PI3K/Akt pathways, which may dominate over canonical signaling and explain the lack of additional ECM induction upon exogenous TGFβ exposure. In diseased states, kidney tubular cells express fibroblast markers, initiating a process in which they lose their epithelial phenotype and acquire new characteristic features of mesenchyme, accompanied by the production of profibrotic factors [48]. Our results showed the nephron segments ephithelium to be present, but our transcriptomic analysis showed that both mutant organoid lines upregulate proximal and glomerular markers while downregulating distal segments. Additionally, *NPHP1^ko1^* displayed reduced *SOX2, PAX2, LIM1,* and *SIX2,* whereas *NPHP1^ko2^* showed upregulation of *SOX2, PAX2,* and *LIM1* with reduced *SIX2* expression. This indicates potential impairments in nephron epithelium maturity, which was confirmed by the detection of bud-shaped and non-contiguous proximal and distal tubules in both *NPHP1^-/-^*organoid lines. Despite we did not explore this further in the present manuscript, the impairment of the nephron epithelium shape and overall structure could additionally be explained by the loss of adherens junctions, of which nephrocystin-1 is a component of. Despite this variability in nephron progenitor dynamics, both *NPHP1*-depleted kidney organoids showed an overall decrease in distal segments epithelium volume, supported by a decrease in gene expression of key markers of these segments, potentially indicating the occurrence of EMT. Supporting this hypothesis, loss of NPH-associated proteins in mice has caused NPH through EMT, driven by the expression of EMT specific genes [49].

To date, the lack in understanding the pathophysiological mechanisms behind the nephrocystin-1 mutations that cause NPH directly limits the development of therapeutic interventions for NPH patients. Currently, the only treatment options for NPH patients are kidney replacement therapy by either dialysis or kidney transplantation. A better understanding of the mechanisms leading to NPH aids in the development of novel drugs that prevent or slow down disease progression. A recent study identified prostaglandin E2 receptors and their downstream pathways, including ECM regulation, to be altered in *NPHP1*-mutants *in vitro* and in animal models, offering a potential therapeutic approach for NPH by E2 agonist treatment [50]. Our RNA-sequencing data revealed significant differences in the prostaglandin E2 receptors *PTGER2*, *PTGER3*, and *PTGER4* in *NPHP1*-mutants (Figure S14, Table S12), as well as altered ECM-receptor pathway, including the impaired expression of collagens, laminins, and integrins upon nephrocystin-1 loss (Figures S15-S16), confirming the phenotype found previously [50] and demonstrating the robustness of our models.

## 4 CONCLUSIONS

Renal ciliopathy-associated proteinshave been linked to DNA stress and/or repair pathways as a disease mechanism in NPH [21, 51–53]. Further investigations of the DDR pathways to better understand the pathophysiology of NPH and eventually develop new treatments for the patients are needed. The present study is the first comprehensive analysis of *NPHP1* and it’s relation to DNA damage pathways in the context of NPH. Our preliminary results blocking the cell cycle to allow time for DNA damage correction, show the potential for targeting the DDR pathways as NPH treatment when it comes to preventing, reverting and/or slowing down DNA damage accumulation due to nephrocystin-1 loss. Altogether, our results further advance our understanding of the molecular mechanism underlying NPH, by providing a robust kidney organoid *NPHP1*-depleted model. Furthermore, our findings pave the way towards identifying new drug targets for treatment of NPH, which point towards DNA damage repair.

Kidney biopsies are not routinely performed in NPHP1 patients due to the invasive nature of the procedure and the fact that diagnosis is usually achieved by genetic testing, and the most appropriate in vivo comparator for our developmental model would be fetal kidney tissue from NPHP1 cases, which is not available to us. Future translational studies about NPHP1 could employ kidney tubuloids or patient-derived iPSC (e.g. from skin biopsies) kidney organoids, which would require biobank ethical permission and infrastructure.

## 5 MATERIALS AND METHODS

### 5.1 Antibodies and reagents

All reagents used were obtained from Thermofisher (Breda, the Netherlands) unless specified otherwise. Unless specified otherwise, primary and secondary antibodies were added at a final dilution of 1:300 and 1:400, respectively. The primary antibody used for the detection of nephrocystin-1 protein was rabbit anti-NPHP1 (#PA5-93140, Invitrogen, Carlsbad, CA, USA). The primary antibodies and probes used for the visualization of the four cell populations of the kidney were rabbit anti-GATA-3 (#5852S; Cell Signaling Technologies, Leiden, The Netherlands), sheep anti-nephrin (#AF4269-SP, R&D systems, Minneapolis, USA), mouse anti-E-Cadherin (#610181, BD Biosciences, Vianen, The Netherlands), and Biotinylated Lotus Tetragonobulus Lectin (LTL, #B-1325, Vector Lab, Brunschwig Chemie, Amsterdam, NL). The primary antibodies against UVC-induced lesions were mouse anti-64PP (#CAC-NM-DND-002, CosmoBio, USA) and mouse anti-CPD (#CAC-NM-DND-001, CosmoBio, USA). Lastly, the primary antibodies used for the assessment of fibrosis were rabbit anti-collagen I (#ab34719, Abcam, Amsterdam, The Netherlands) and sheep anti-fibronectin (#AF1918, R&D systems, Dublin, Ireland, 1:400). TGFβ was added at a final concentration of 50ng/mL for 24 h. The primary antibody for the detection of cleaved caspases 3 was rabbit anti-cleaved caspases 3 (#PA5-114687). The detection kit used for the assessment of senescence was the Cell Event^TM^ Senescence green Detection kit (#C10850, Invitrogen, 1:1000). For the secondary fluorophore conjugation, we used Alexa-647 Donkey anti-sheep (#A-21206), Alexa-568 Donkey anti-rabbit (#A10042), Alexa-488 Donkey anti-rabbit (#A-21206), Alexa-488 Donkey anti-mouse (#A21202, and Alexa-405 streptavidin conjugate (#S32351).

### 5.2 Cell lines and iPSC culture

The iPSC line used in the current study (iPS134Cl2; ID: CMO 2015-1543) were generated from human erythroblasts from a healthy pediatric caucasian patient using the Yamanaka factors by the Stem Cells Technology Center at Radboud University Medical Center (SCTC, Radboud UMC, Nijmegen, The Netherlands) [54]. Two different knockout lines originated from the same donor iPSC line were generated for this study. Their genomic status was assessed with Sanger Sequencing once after clonal expansion and in several other instances throughout the project to ensure no random genomic corrections/alterations in the mutant region. The *NPHP1^-/-^*compound heterozygous line harbors a compound heterozygous frameshift mutation in *NPHP1* (allele 1: c.287_288insT (predicted to introduce a premature stop codon), allele 2: c.288_288delG (predicted to introduce a premature stop codon) and is referred in the main text as *NPHP1*^ko1^. The *NPHP1^-/-^* homozygous line harbors a homozygous frameshift mutation in *NPHP1* (allele 1: c.287_288insT, allele 2: c.287_288insT (predicted to introduce a premature stop codon) and is referred in the text as *NPHP1*^ko2^. To exclude the possibility that genetic anomalies arising during the gene editing process could account for phenotypic differences, we performed karyotyping on all cell lines (Supplementary File 1). No chromosomal abnormalities were detected in either line, indicating that the divergent phenotypes are not attributable to gene-editing-induced or culture-induced genetic anomalies.

The iPSCs were cultured in Essential 8 (E8) medium containing E8 supplement and 100 μg/mL Penicillin-Streptomycin (E8 complete medium), all purchased from GIBCO (Life Technologies, Paisley, United Kingdom) in well 6 well plates that were coated with 1% Geltrex (ThermoFischer Scientific, Breda, NL). Cells were cultured at 37°C in a humified atmosphere in the presence of 5% CO_2_ and passaged at a split ratio of 1:3/1:6 every 3 to 5 days, respectively. All iPSC lines were frozen in E8 culture media with 10% DMSO. Cells were used for experiments at passages ranging from 20 up to 39, keeping the passage number amongst all NPHP1 lines within similar range for experimental purposes. The *NPHP1*^WT^ iPSCs were cryopreserved at passages 11 to 40 while *NPHP1* mutant iPSC lines were cryopreserved at passages 20 to 40. Mycoplasma testing was performed routinely on a monthly basis and stock cultures were evaluated for bacterial, yeast, and fungal infections every 2-3 days.

### 5.3 Kidney organoids differentiation from iPSCs

For the differentiation into kidney organoids, iPSCs were first washed in PBS and resuspended in TrypleSelect for dissociation. After centrifugation and resuspension in Essential 8 medium, cells were seeded on Geltrex-coated 6-well plates with a density of 250.000 cells/well. The differentiation procedure is an established protocol by Jansen et al. (2022) [55] from the original protocol from Takasato et al. (2015) [56], which we have previously published [57]. In short, iPSCs were differentiated into progenitor kidney cells during 7 days in 2D in E6 medium using CHIR-99021 (6μM, #4423, R&D Systems), fibroblast growth factor 9 (FGF9, #130-110-920, 200 ng/ml, R&D Systems) and heparin (1 μg/ml, Sigma-Aldrich, Zwijndrecht, Netherlands). From day 3, half of the wells were switched to E6 medium containing FGF9 and 1 µg/ml heparin and maintained in these conditions until day 7 to generate anterior intermediate mesoderm. The remaining wells stayed in E6 medium with CHIR99021 until day 5 to induce posterior intermediate mesoderm. From day 5 onward, all wells were cultured in E6 medium supplemented with FGF9 and heparin. On day 7, cells were aggregated and seeded on 6-well inserts (CellQart #9300402, Northeim, DE) and cultured in an air-liquid interface. The medium was replaced with E6 supplemented with FGF9 and heparin for an additional 5 days. After that, the organoids were cultivated in E6 supplemented with human epidermal growth factor (hEGF, 10 ng/ml, Sigma-Aldrich) and bone morphogenetic protein-7 (BMP7, #130-093-818, 50 ng/ml, R&D Systems) until they were harvested on day 7+18. Medium of organoids was refreshed every 2 days, or every 3 days during weekends.

### 5.4 Nucleofection of iPSCs

The iPSCs were nucleofected with a 2b-Nucleofector (Lonza, Basel, Switzerland). Cells were preincubated for 1 h with 1X RevitaCell (#A2644501). Cell pellets were collected after washing the wells twice with PBS and by detaching the cells at 37 °C for 2 min with TripLE Select Enzyme (1X, no phenol red), after which the well was neutralized with E8 medium and centrifuged for 3 min at 400 rcf. The pellets were adjusted to 1 million cells per condition and diluted with complete Nucleofector^TM^ Solution V (Lonza). For the gRNA duplex assembly, 2000 μM of crRNA (5’ AAGCUUACCCAACAACUGCA3’) was mixed with 2000 μM of tracrRNA at 95°C for 5 min and, subsequently, 61 μM of Cas9 was added to produce the ribonucleoprotein complex (RNP). After 20 min, 5.3 μl of Cas9 Electroporation Enhancer (IDT) was added, and the diluted iPSCs were nucleofected with the previous mixture using the program B-016, following the manufacturer’s instructions. The cells were re-plated and cultured in E8 culture medium supplemented with 1X RevitaCell in a 6-well plate (Greiner Bio-one) for 24 h at 37 °C.

### 5.5 Fluorescence-activated Cell Sorting (FACS)

Cells were dissociated into single cells 16-24 h after nucleofection, as described in the section above Collected single cells were resuspended in E8 complete medium and transferred through a 40 μm cell strainer (Corning Incorporated, NY, USA) to obtain single cells. Next, cells were sorted into single cells at the Flow Cytometry and Cell Sorting Facility at the Faculty of Veterinary Medicine (Utrecht, the Netherlands). Cells that showed fluorescence in the 488 or 550 (depending on the used tracRNA) spectra were selected and seeded as single cell per well in a Geltrex coated 96-well plate containing E8 completed medium supplemented with Clone-R (STEMCELL Technologies, Vancouver, Canada). Plates were spun down to increase single cell attachment and were placed at 37 °C in a humified atmosphere in the presence of 5% v/v CO_2_ until colonies were formed (∼2-3 weeks after sorting), refreshing the media every 3-5 days.

### 5.6 DNA extraction and PCR

The iPSCs were gathered as single cells as described in “Cell lines and iPSC culture”. Cell pellets were washed once with PBS and the DNA was extracted using the QIAamp DNA Mini Kit according to the manufacturer’s instructions (Qiagen, Venlo, the Netherlands). Polymerase chain reaction (PCR) was performed at 60 °C with 200 ng of isolated DNA and using 5’ TGCATTCCAACACAATTGCCT3’ and 5’ CCTGAGGGCAGAGACTATGAC3’ as forwards and reverse primers, respectively. The PCR product was purified with a PCR Purification Kit (Qiagen, Hilden, Germany) and sent for Sanger-sequencing to Macrogen (Amsterdam, the Netherlands). The knock-out efficiency was analyzed using TIDE web tool (https://tide.nki.nl/) and the ICE software was used for Sanger-Sequencing traces visualization (Synthego Performance Analysis, ICE Analysis. 2022. v3.0, https://ice.synthego.com).

### 5.7 In-silico prediction analysis

The DNA variant effect prediction tool MutationTaster (http://www.mutationtaster.org/) was used to predict the functional consequences of the INDELs in the *NPHP1* gene and I-TASSER (http://zhanglab.ccmb.med.umich.edu/I-TASSER/) [58] was used to model the 3D structure of both full-length and mutated nephrocystin-1 proteins caused by the biallelic homozygous and compound heterozygous in *NPHP1* generated by CRISPR/Cas9. Starting from the predicted 3D model, the COFACTOR webserver (https://zhanggroup.org/COFACTOR2) [59] was used to identify potential protein-ligand binding interactions by using local and global structure matches. To further examine possible binding partners of nephrocystin-1, STRING (Search Tool for the Retrieval of Interacting Genes/Proteins) and PEPPI (Pipeline for the Extraction of Predicted Protein-Protein Interactions) were used. STRING is a protein-binding prediction tool that integrates data from experimental studies, computational predictions, and text-mining approaches. It provides information about protein-protein interactions and functional associations. Confidence scores are assigned to interactions, and the results are visualized as interactive networks. PEPPI is a computational program for predicting protein-protein interactions (PPI). It estimates the likelihood of direct interactions between a pair of protein amino acid sequences using multiple methods: structural homology via multimeric threading, sequence homology through BLAST search using experimental data, functional association from the STRING database, and machine learning-based classification. The final likelihood ratio, derived from a naive Bayesian consensus model, expresses the probability of interaction relative to non-interaction.

### 5.8 Bulk RNA sequencing and analysis

For RNA-Sequencing analysis, a single kidney organoid was used for every biological replicate. The kidney organoids underwent two washes with ice-cold PBS, followed by dissociation in 350 µL of cold lysis buffer (RLT Buffer, QIAGEN). Subsequently, the samples were stored at -80°C until submission to the USEQ Utrecht Sequencing Facility (Utrecht, the Netherlands). For each sample, three biological replicates were submitted for sequencing. For two samples (*NPHP1*^ko1^ Untreated and *NPHP*^WT^ +UVC) the RNA quality was suboptimal and therefore these replicates were not included in the analysis. The raw transcript counts provided were processed using the web application IDEP versions 1.12-2.0 [60] to generate all the plots presented in the results section of this manuscript. In short, the raw data (total read counts) were imported into the IDEP1.12 web tool, normalized using counts per million (CPM), and pre-processed by filtering out genes with less than 20 CPM (which allowed for the retention of 17.1% of the transcripts). Subsequently, the raw data were transformed for clustering using the Edge:log2 (CPM + c) method. The principal component analysis (PCA) were constructed from the linear transformation of the data, such that the first components point to the direction of the most variation among the samples. The differential gene expression between the experimental groups (DEG1 and DEG2) was analyzed employing the DeSeq2 method, with a false discovery rate (FDR) cutoff of 0.05 and a minimum fold-change of 1.5. From the identified DEG1 and DEG2, the top enriched pathways were obtained, and subsequent pathway trees and networks were generated. For the pathway analysis, absolute values of fold changes were utilized with the GAGE method, sorting pathways by biological process, and setting the FDR-cutoff at 0.2. For the generation of the DDR pathway-specific heatmaps, we selected the best known DDR-related pathways and plotted the Z-scores from the CPM values, and plotted the results using GraphPad 9.0 (GraphPad software, La Jolla, CA, USA). Finally, the KEGG-database and Pathview were employed to integrate up- and down-regulated genes within each available KEGG pathway. Access to the raw RNA sequencing data is available upon request. In the RNA-Seq supplementary and raw files, *NPHP1*^ko1^ appears as “NPHP1_C1” and *NPHP1*^ko2^ as “NPHP1_C3”. Additionally, the raw data were deposited in the public repository Figshare (https://figshare.com).

### 5.9 Immunostainings

After completing the maturation protocol, the organoids were washed 3 times with PBS and fixed with 2% PFA solution (PierceTM 16% formaldehyde (w/v), methanol-free, ThermoFisher) on ice for 20 min. After incubation, PFA was removed, and organoids were washed 3 times with PBS. Next, organoids were used for stainings or organoids were stored in 0.1% v/v formalin at 4°C. For antibody stainings, organoids were blocked in blocking buffer (BB) containing 10% v/v donkey fetal serum (GIBCO) and 0.06% Triton-X (ThermoFisher Scientific) in 1× PBS. Next, primary antibodies in BB were added and incubated overnight while gently shaking at 4°C. After incubation, primary antibody solutions were removed and organoids were washed 3 times with 1× PBS for 10 min while gently shaking at room temperature. Subsequently, secondary antibodies were added in 0.3% v/v triton-X in 1× PBS and incubated for 3 h while gently shaking at room temperature. Next, secondary antibody solutions were removed and organoids were washed 3 times with 1× PBS for 10 min while gently shaking at room temperature. The Alexa Fluor 647 Anti-gamma H2A.X probe was incubated overnight at 4°C with a gentle shaking diluted in 0.3% v/v triton-X in 1× PBS. The Senescent probe staining was performed using the CellEventTM Senescence Green Detection Kit (InvitrogenTM) following the manufacturer’s instructions. Finally, organoids were mounted in Prolong gold containing DAPI. Images were captured using the confocal microscope Leica TCS SP8 X (Leica Biosystems, Amsterdam, The Netherlands).

### 5.10 SDS- Polyacrylamide Gel Electrophoresis (SDS-PAGE) and Western-Blot

Equal amounts (BCA assay normalized) of each sample were diluted in Laemmli Sample Buffer (Bio-Rad) with 100mM DTT and heated at 95 for 5 min. Samples and precision plus pre-stained marker (Bio-Rad) were loaded onto a 4-20% TGX Gel (Bio-Rad) and run under reducing conditions in 1xTris-Glycine-SDS running buffer (Bio-Rad) at 100V for 90 min. Proteins were transferred to PVDF membrane using the Trans-Blot turbo system (Bio-Rad). The membrane was blocked for 1h in 5%Milk/PBS/0,1%Tween-20. NPHP-1 polyclonal antibody (#PA5-93140, Invitrogen, Carlsbad, CA, USA) was added at 4 o/n in 5%Milk/PBS/0,1%Tween-20. The membrane was washed in PBS/0,1%Tween20 and incubated in 5%Milk/PBS/0,1%Tween-20 for 1h at RT with HRP labelled Goat-anti-Rabbit IgG (DAKO). After washing in PBS/0,1%Tween20 the proteins were visualized using ECL Clarity (Bio-Rad) and the CemiDoc Touch Imaging System (Bio-Rad).

### 5.11 Karyotyping

Karyotyping of iPSC lines was performed by standard GTG-banding (G-banding with trypsin and Giemsa) at the Genoomdiagnostiek department (Genetics, UMC Utrecht, Utrecht, the Netherlands), an ISO 15189-accredited clinical diagnostics laboratory. Twenty metaphase spreads were analyzed per line at passage 45 (KO1) and passage 47 (KO2). Both iPSC lines displayed a normal female karyotype (46,XX).

### 5.12 Compound testing

To evaluate whether the prevention of cell division would allow for DNA damage repair, we used RO-3306, a recently described potent selective ATP-competitive inhibitor of CDK1, which prevents mitotic entry and arrests cell division at the G2/M boundary. RO-3306 was dissolved in DMSO and used at a final concentration of 10 nM and added to the kidney organoids for 24 h. The negative control was treated with the same concentration of DMSO as the treated condition for effect normalization.

### 5.13 UVC light exposure

At the end of the maturation protocol (d7+18), organoids were exposed to 25 J/m2 of UVC light using a UVP CL-1000 UV Crosslinker by introducing the plate they were cultured into the crosslinker without the lid. The time of exposure is determined automatically by the crosslinker depending on the UVC dose chosen. Upon UVC treatment, all organoids were kept in the dark in the incubator until harvesting. UVC-induced DNA damage is a rapid occurring phenomenon, we therefore harvested the organoids rapidly after exposure including several timepoints between 15 min up to 48 h, at which they were constantly kept in the dark in the incubator.

### 5.14 Data analysis

Every experiment was performed in biological replicates (N=3), including at least 3 technical replicates each, unless specified otherwise. Each biological replicate represents an independent differentiation batch, originated from different passage numbers of the iPSCs. Results are shown as the mean ± standard error of the mean (SEM) and all p-values <0.05 were considered statistically significant. The threshold of the p-values are stated on each figure legend. All gathered numeric data was analyzed using GraphPad 9.0 (GraphPad software, La Jolla, CA, USA). For statistical analysis, one-way ANOVA was performed followed by Tuckey post-hoc analysis unless specified otherwise. In some cases, two-way ANOVA followed by a Dunnett or Šidák correction for multiple comparisons. Image analysis was performed with ImageJ (ImageJ software version 2.9.0/1.53t, National Institutes of Health, Bethesda, MD, USA). For the semi-quantification of 64PPs, CPDs, and yH2Ax in Figure 5, Figure 6, and Figure S4, we adopted a targeted imaging approach. During acquisition, we first used DAPI staining to identify tubular structures and then zoomed into these regions for high-resolution (63× + 5× digital) imaging. We acknowledge that with DAPI staining alone it is not possible to distinguish between proximal and distal tubules. To account for this, we ensured representative sampling by imaging three different organoids per condition, and within each organoid, capturing three independent tubular regions. While this approach does not allow precise nephron segment identification, it maximizes the likelihood of capturing multiple proximal tubules and provides a robust assessment of photolesion distribution and yH2Ax presence within tubular structures. For semi-quantitative analysis of fluorescent images, RGB images were converted to 8-bit. Subsequently, grey scale thresholds were adjusted to only represent true staining. Pixel area limited to the threshold was measured for the different stainings.Eventually, the pixel area of stainings was divided by the pixel area of DAPI for each image for normalization.

## 6. RESOURCE AVAILABILITY

### Lead Contact

Further information and requests for resources and reagents should be directed to and will be fulfilled by the lead contact, Anne Metje van Genderen.

### Materials availability

All unique/stable organoid models generated in this study are available from the lead contact with a completed materials transfer agreement and only after consultation and approval of the Radboudumc Stem Cell Technology and differentiation Center.

### Data and code availability

Bulk RNA-seq raw data is deposited at the public repository Figshare and will become publicly available as of the date of publication (figshare.com; file ID: 54147335). Microscopy data reported in this paper will be shared by the lead contact upon request. Any additional information required to analyze the data reported in this paper is available from the lead contact upon request. All collected data was managed and stored following the FAIR principles.

## Acknowledgements

The authors wish to thank the Radboudumc Stem Cell Technology and differentiation Center (https://www.radboudumc.nl/en/research/radboud-technology-centers/stem-cells) for reprogramming and characterizing the cell lines. The authors thank Alinda Berends and Bart Blokhuis or helping with the rebuttal experiments.

## Author contributions

Conceptualization, E.S.G., N.V.A.M.K., A.M.E., R.M., G.G.S., A.M.G. and M.J.J.; methodology, E.S.G. and S.B. ; investigation, E.S.G. and S.B. ; visualization, E.S.G. G.G.S. and A.M.G ; funding acquisition, M.J.J. and R.M. ; project administration, A.M.G., M.J.J. and R.M. ; supervision, R.M., G.G.S., A.M.G. and M.J.J. ; writing – original draft, E.S.G. and S.B.

## 8. SUPPLEMENTARY MATERIALS

### 8.1 Supplementary Figures (S1-S9)

**Figure S1.**
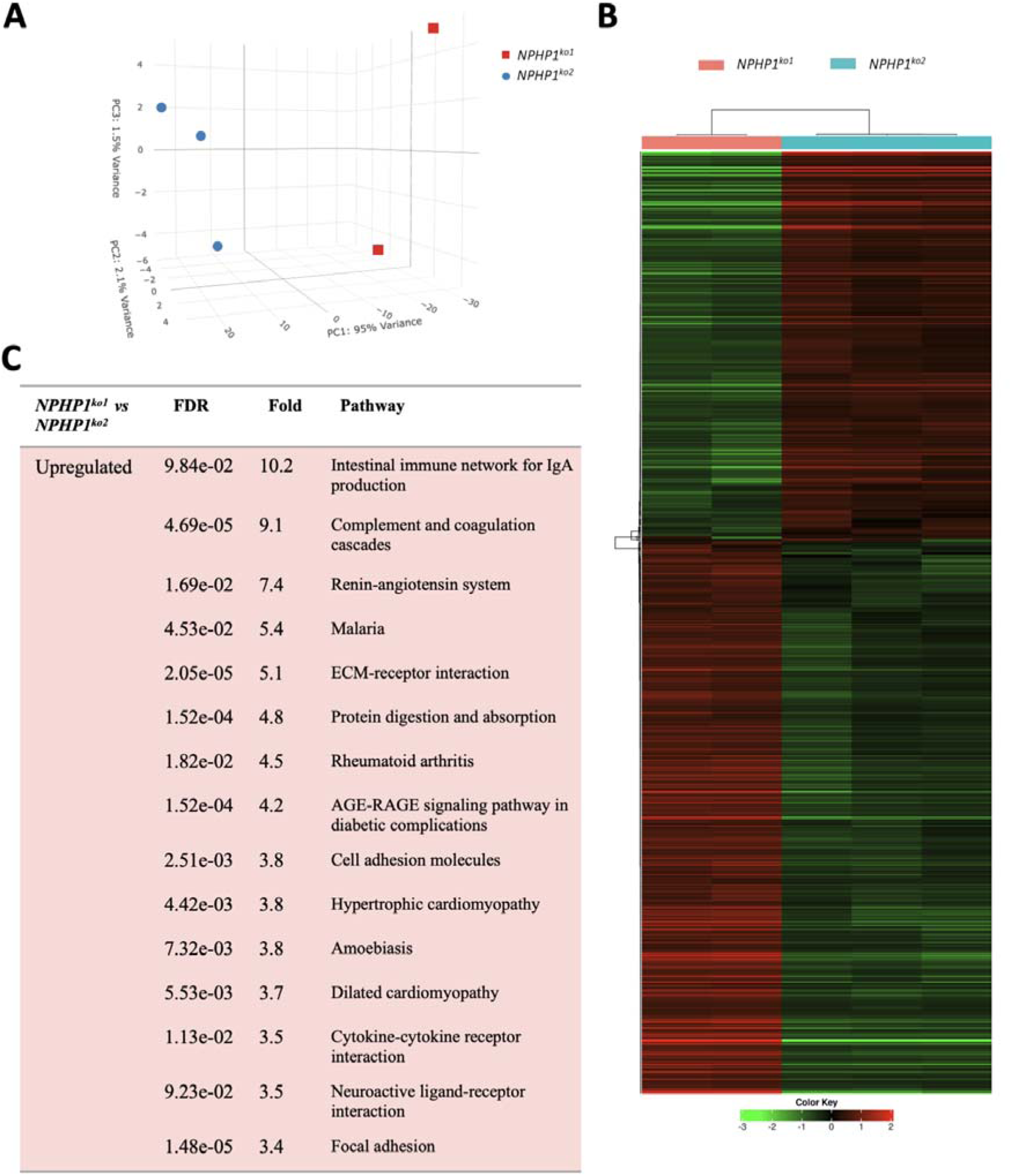
Comparative transcriptomic analysis between *NPHP1^ko1^* and *NPHP1^ko2^*. **(A)** Principal component analysis (PCA) results represented in a 3-dimensional plot. **(B)** Heatmap representation of the unsupervised clustering of the top 1000 genes. **(C)** Table representing the top 15 pathways after DEG enrichment analysis when comparing *NPHP1^ko^*^1^ to *NPHP1^ko2^*, shown in descending order based on their fold-change. All 15 top pathways were upregulated. N=3 (except for *NPHP1^ko^*^1^, for which N=2). Each biological replicate represents a single kidney organoid of a different differentiation round.

**Figure S2.**
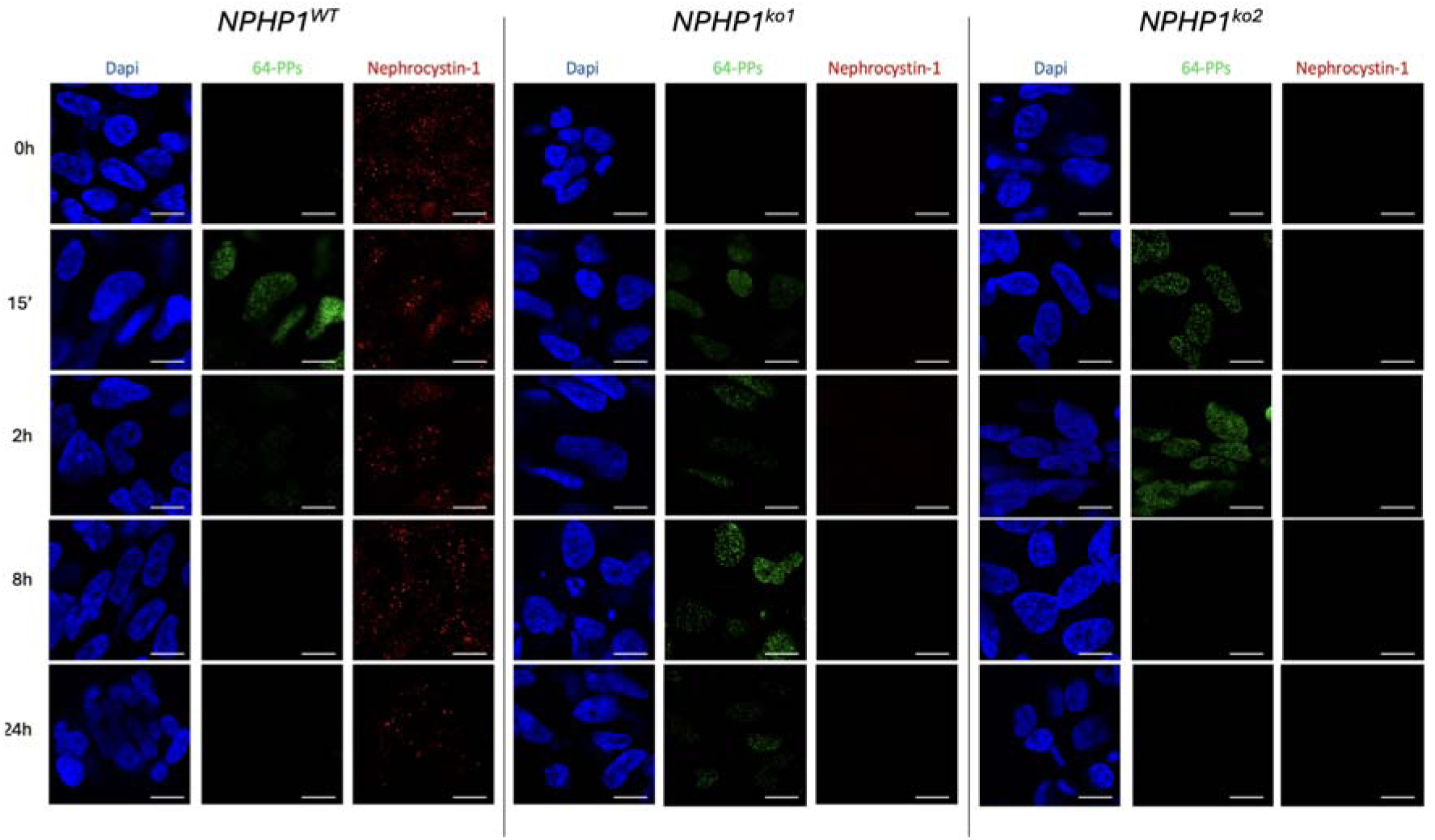
Unmerged images from Figure 5A in the main manuscript. Representative immunostainings at 63X plus 5X digital zoom of 64PPs in addition to nephrocystin-1 in all three organoid lines at different timepoints after exposure to 25 J/m2 of UVC light.

**Figure S3.**
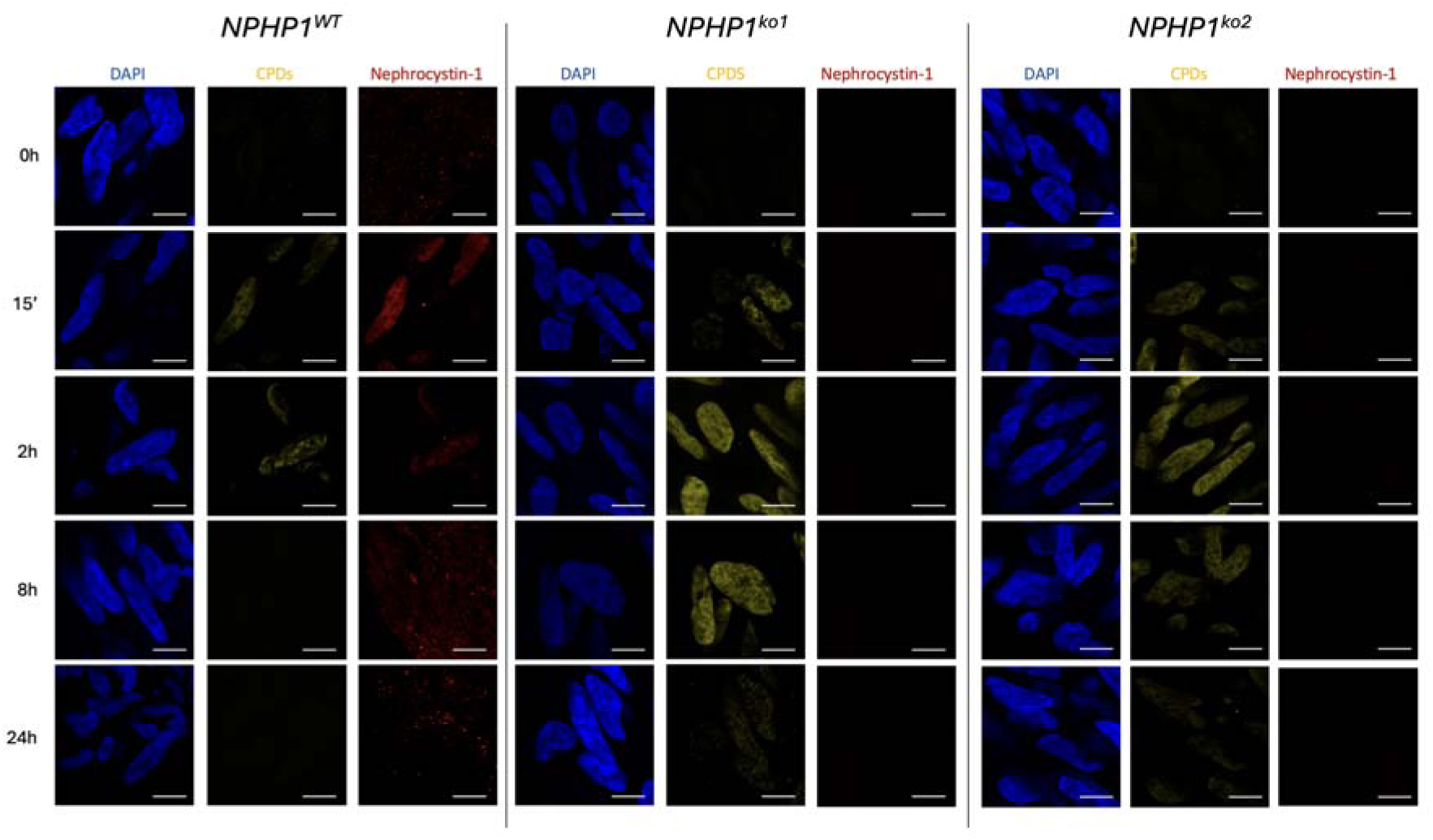
Unmerged images from Figure 5B in the main manuscript. Representative immunostainings at 63X plus 5X digital zoom of CPDs in addition to nephrocystin-1 in all three organoid lines at different timepoints after exposure to 25 J/m2 of UVC light.

**Figure S4.**
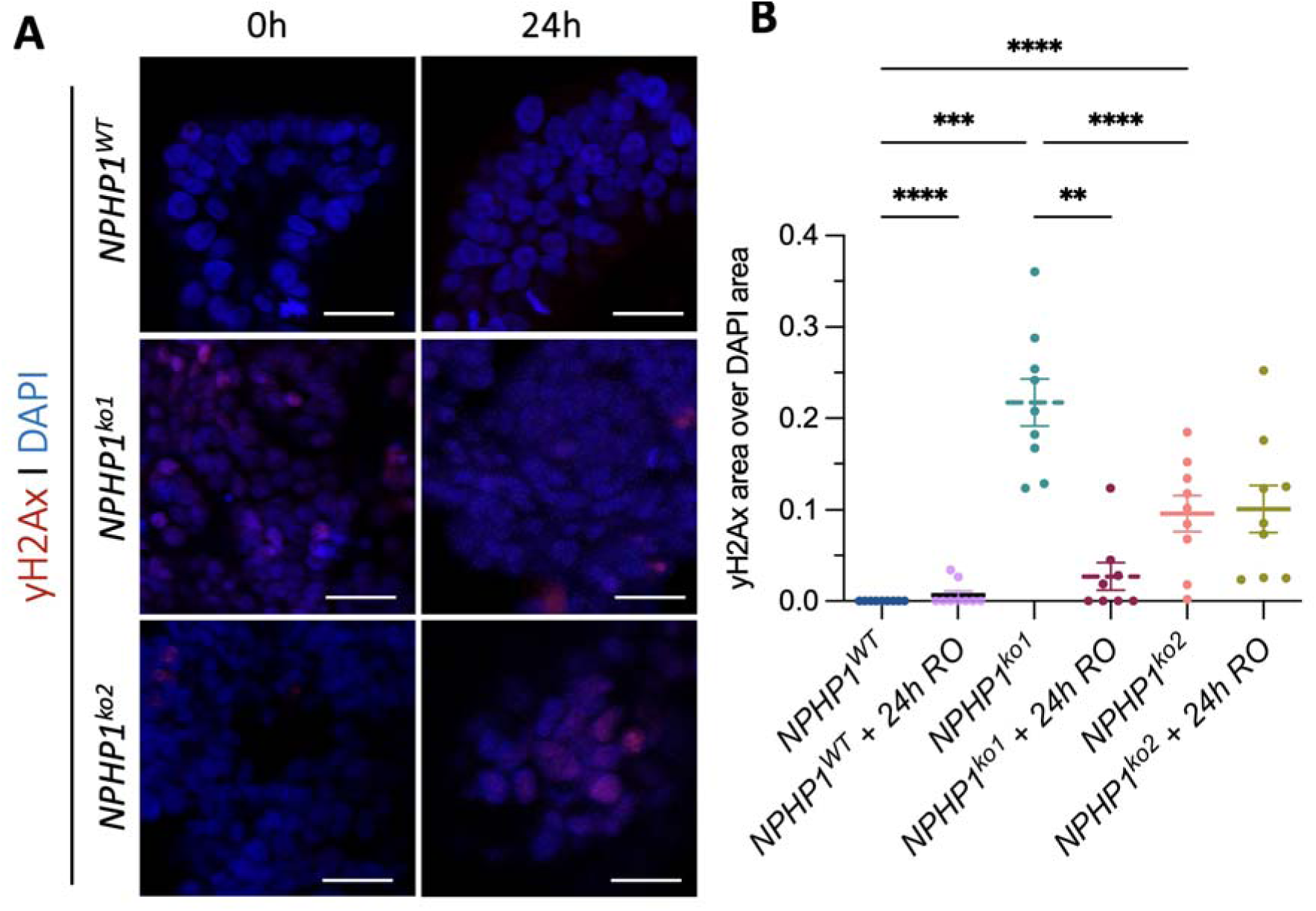
Treatment with RO-3306 ameliorates DNA damage signalling in *NPHP1^ko^*^1^. **(A)** Representative immunofluorescence images at 63X plus 3X digital zoom of all three organoids under untreated conditions and after 24 h of treatment with 10 nM RO-3306. **(B)** Image quantification of the immunofluorescent staining calculated as area of gH2AX over the total DAPI area. A one-way ANOVA analysis was performed (N=3 biological replicates, in which each represents one kidney organoid of a different differentiation round. Within each biological replicate, each technical replicate represents a different region of the same kidney organoid ()p<0.05). Scale bar = 20 µm.

**Figure S5.**
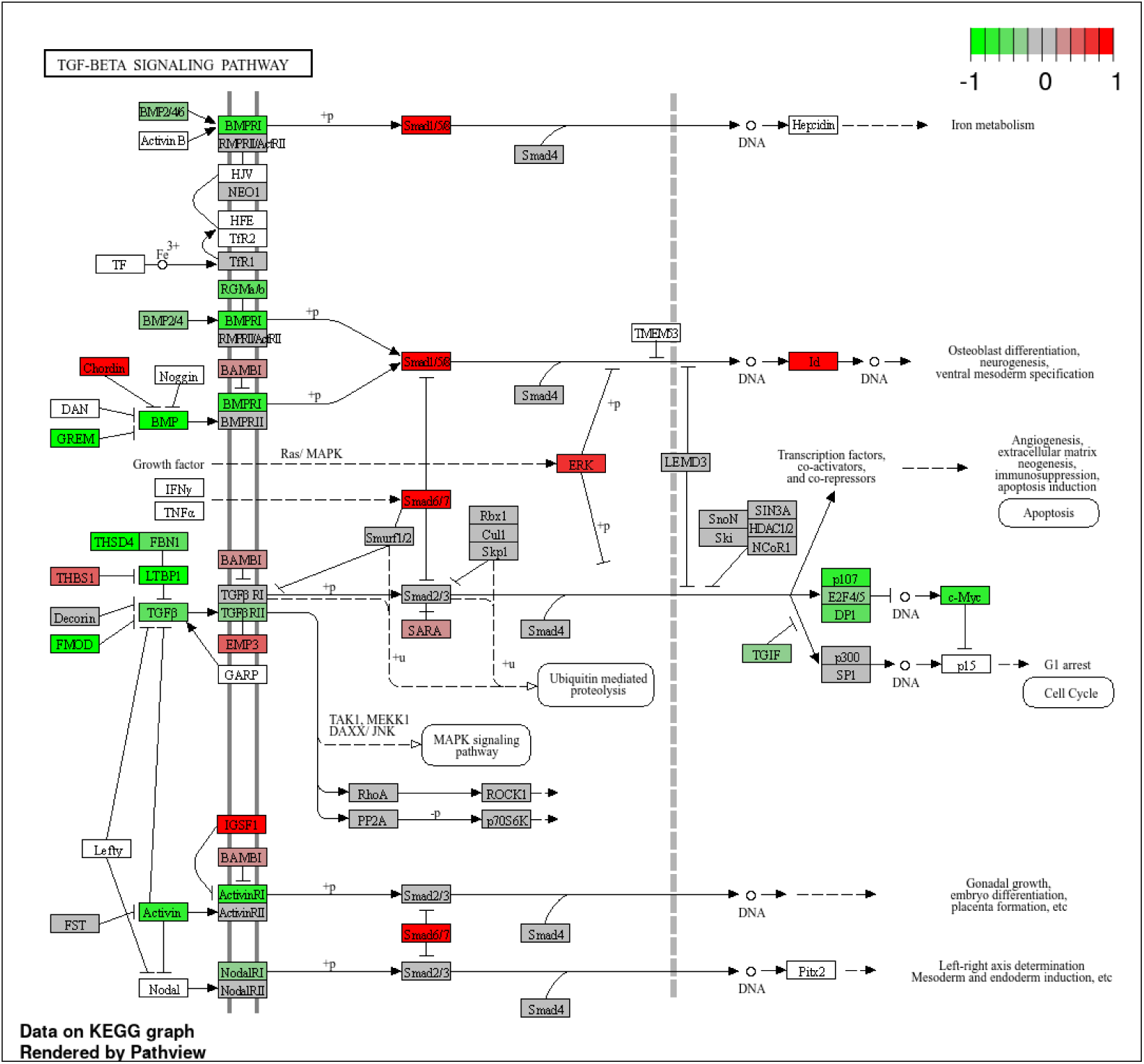
Schematic representation of the TGFβ signaling pathway (KEGG ID: hsa04350) using KEGG pathways analysis for *NPHP1^ko1^* kidney organoids compared to *NPHP1^WT^*.

**Figure S6.**
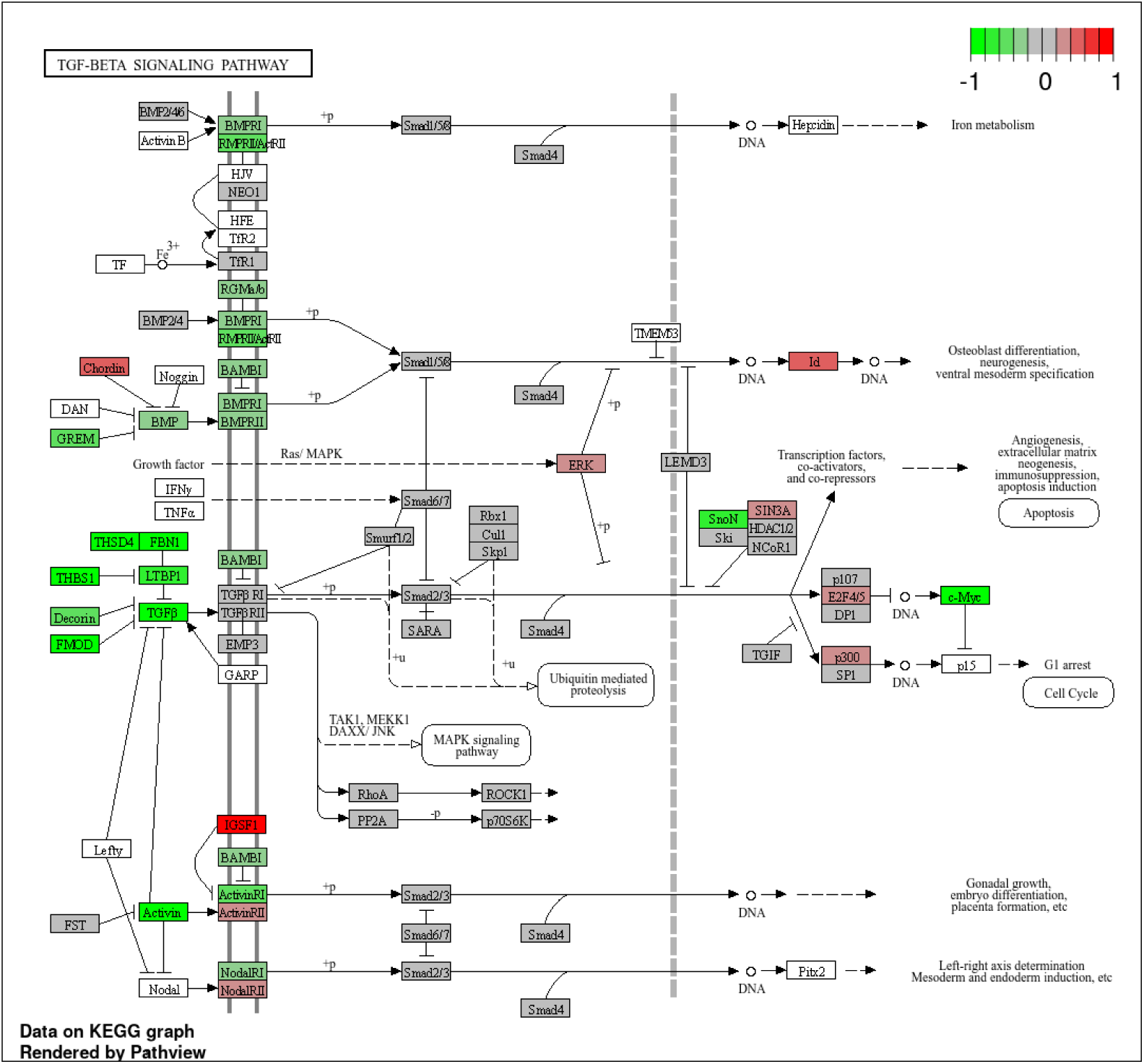
Schematic representation of the TGFβ signaling pathway (KEGG ID: hsa04350) using KEGG pathways analysis for *NPHP1^ko2^* kidney organoids compared to *NPHP1^WT^*.

**Figure S7.**
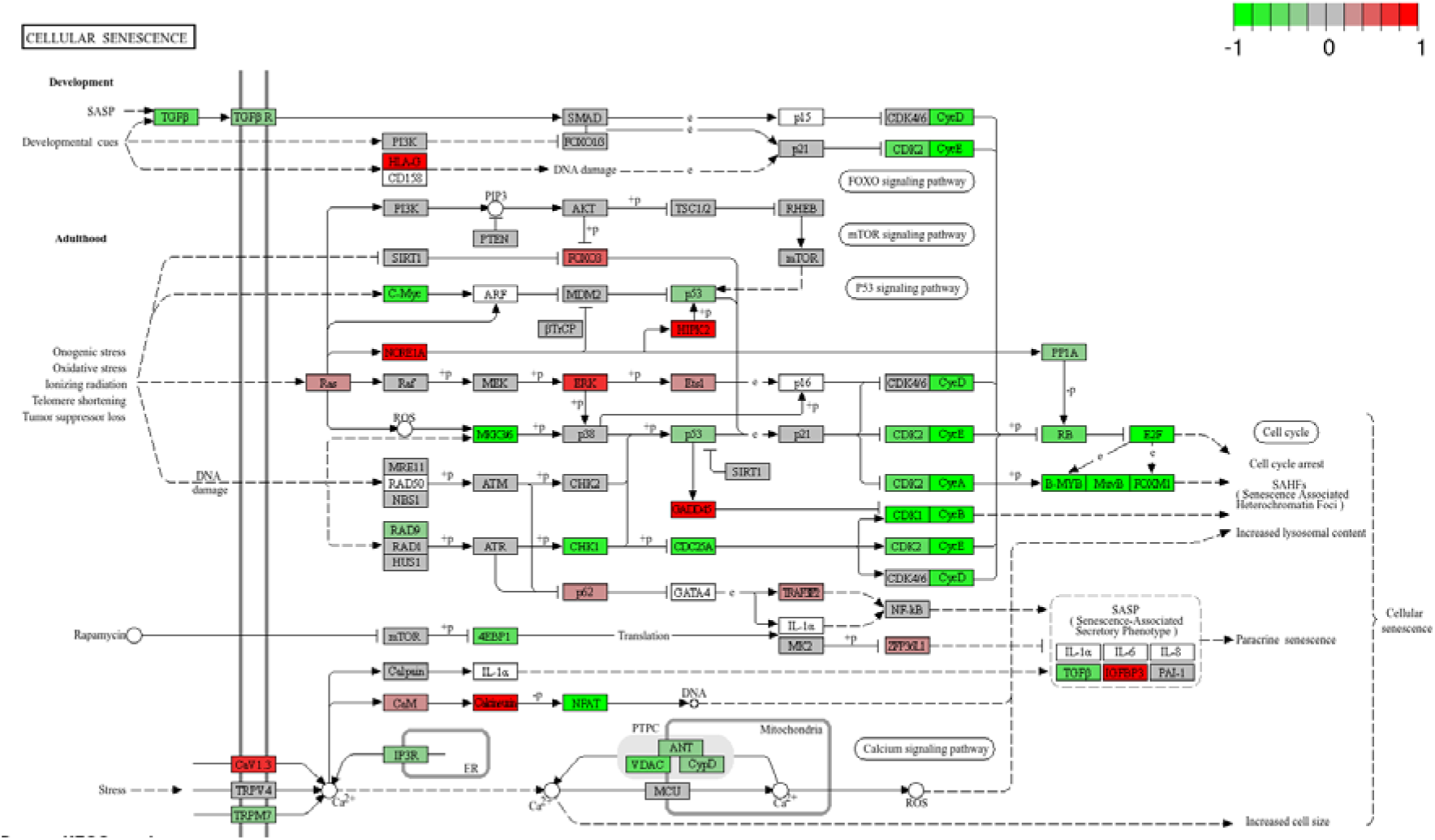
Schematic representation of the cellular senescence pathway using KEGG pathways analysis for *NPHP1^ko1^* kidney organoids compared to *NPHP1^WT^*.

**Figure S8.**
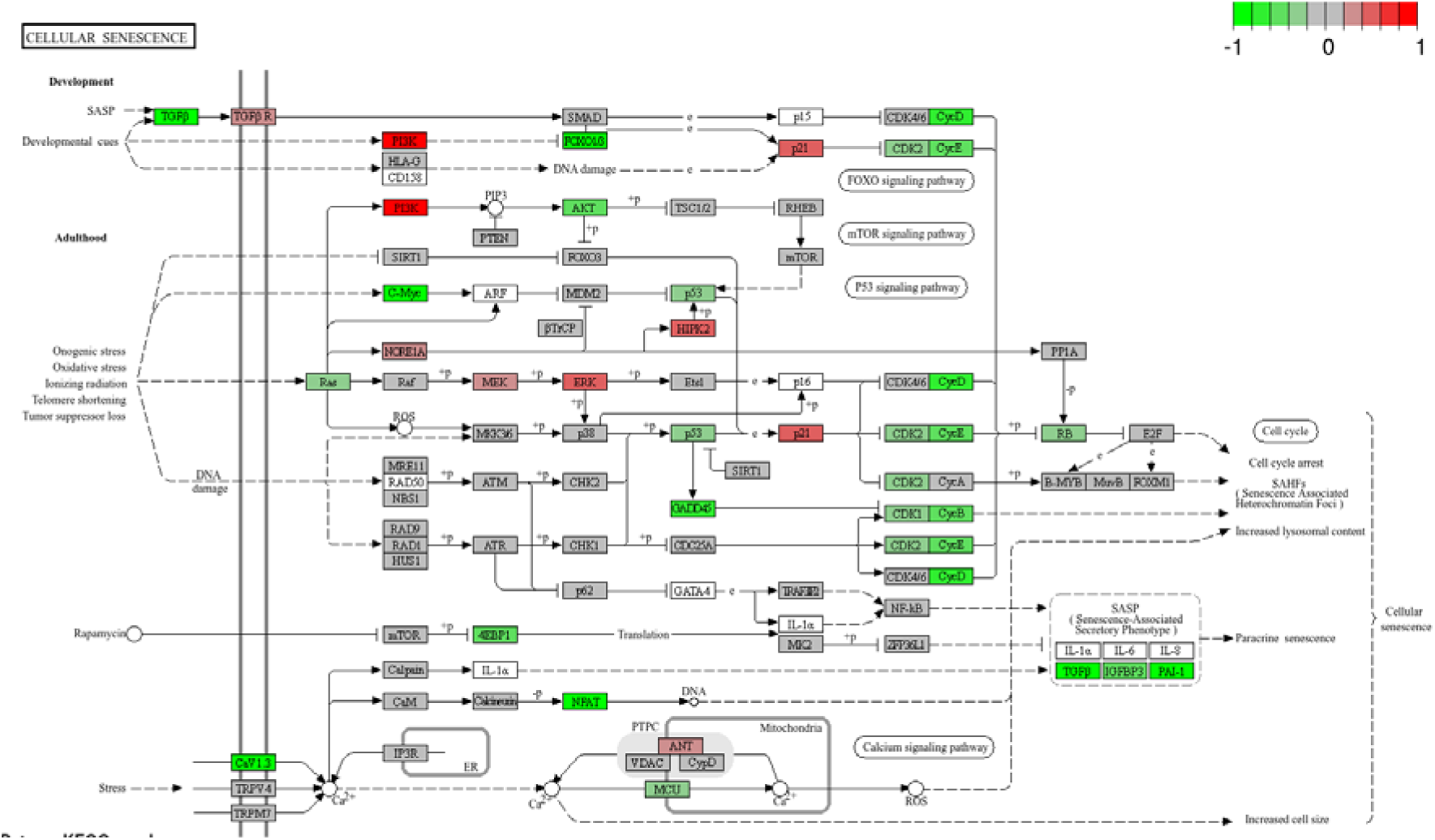
Schematic representation of the cellular senescence pathway using KEGG pathways analysis for *NPHP1^ko2^* kidney organoids compared to *NPHP1^WT^*.

**Figure S9.**
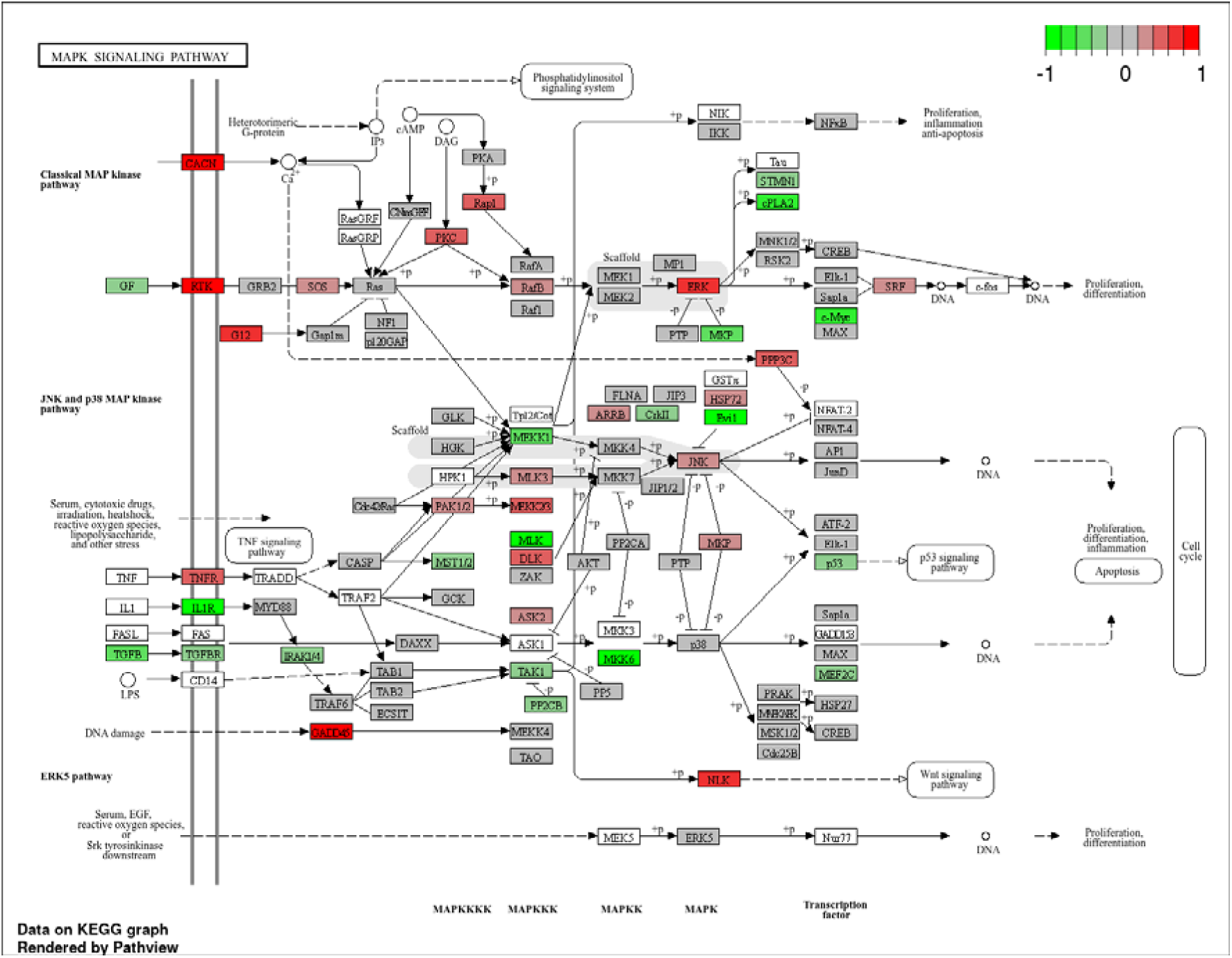
Schematic representation of the MAPK signaling pathways using KEGG pathways analysis for *NPHP1^ko1^* kidney organoids compared to *NPHP1^WT^*.

**Figure S10.**
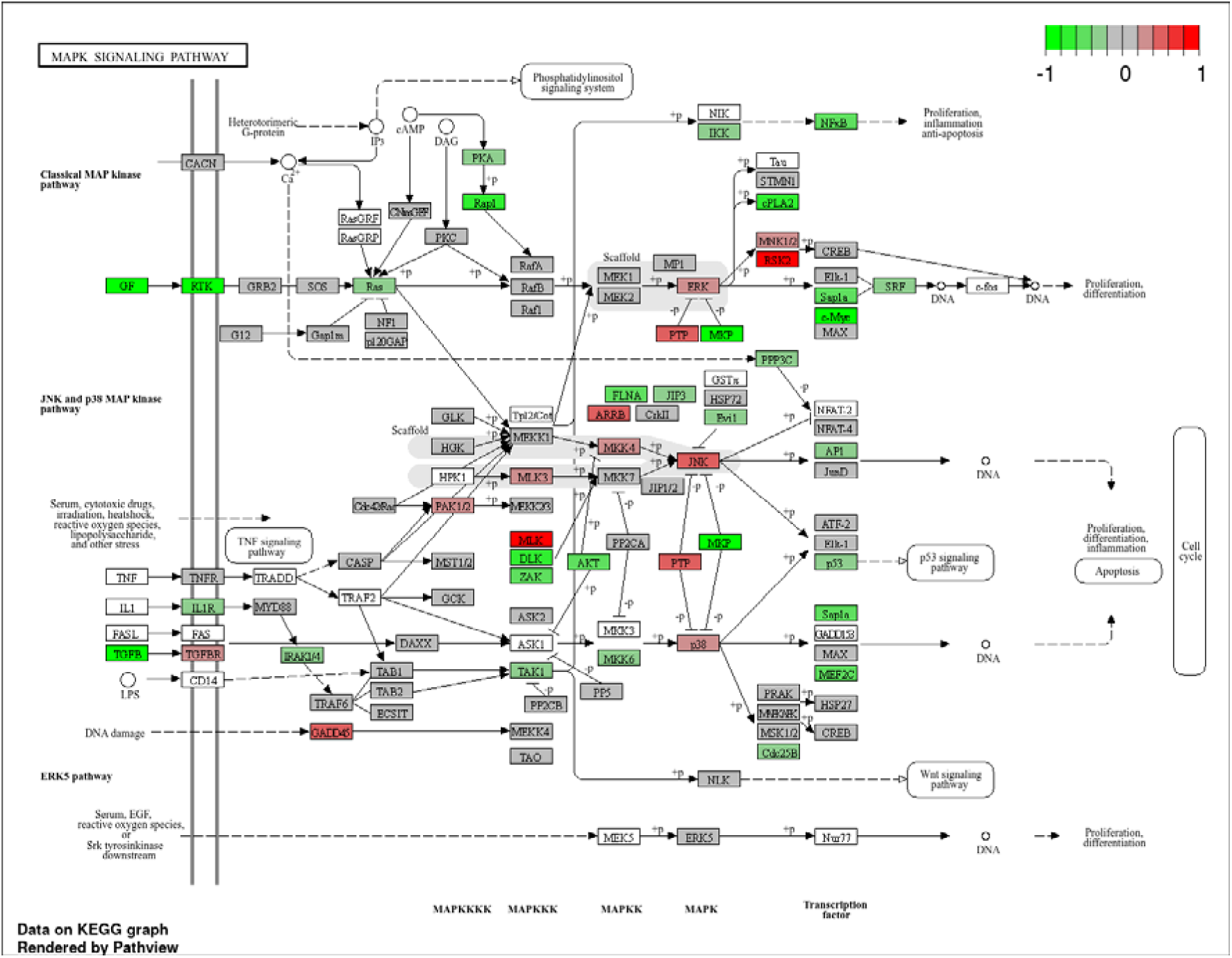
Schematic representation of the MAPK signaling pathways using KEGG pathways analysis for *NPHP1^ko2^* kidney organoids compared to *NPHP1^WT^*.

**Figure S11.**
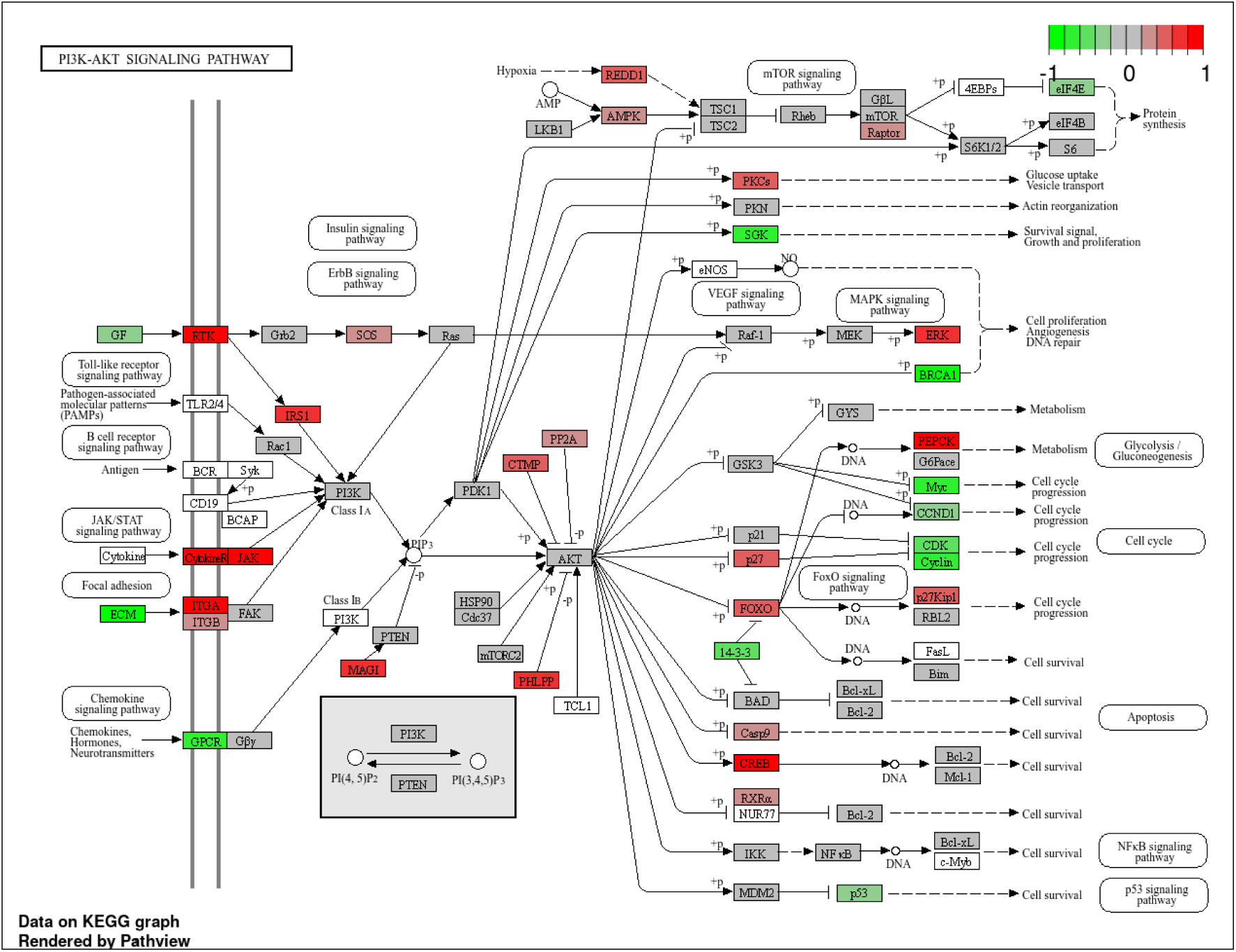
Schematic representation of the PI3K-AKT signaling pathway signaling pathways using KEGG pathways analysis for *NPHP1^ko1^* kidney organoids compared to *NPHP1^WT^*.

**Figure S12.**
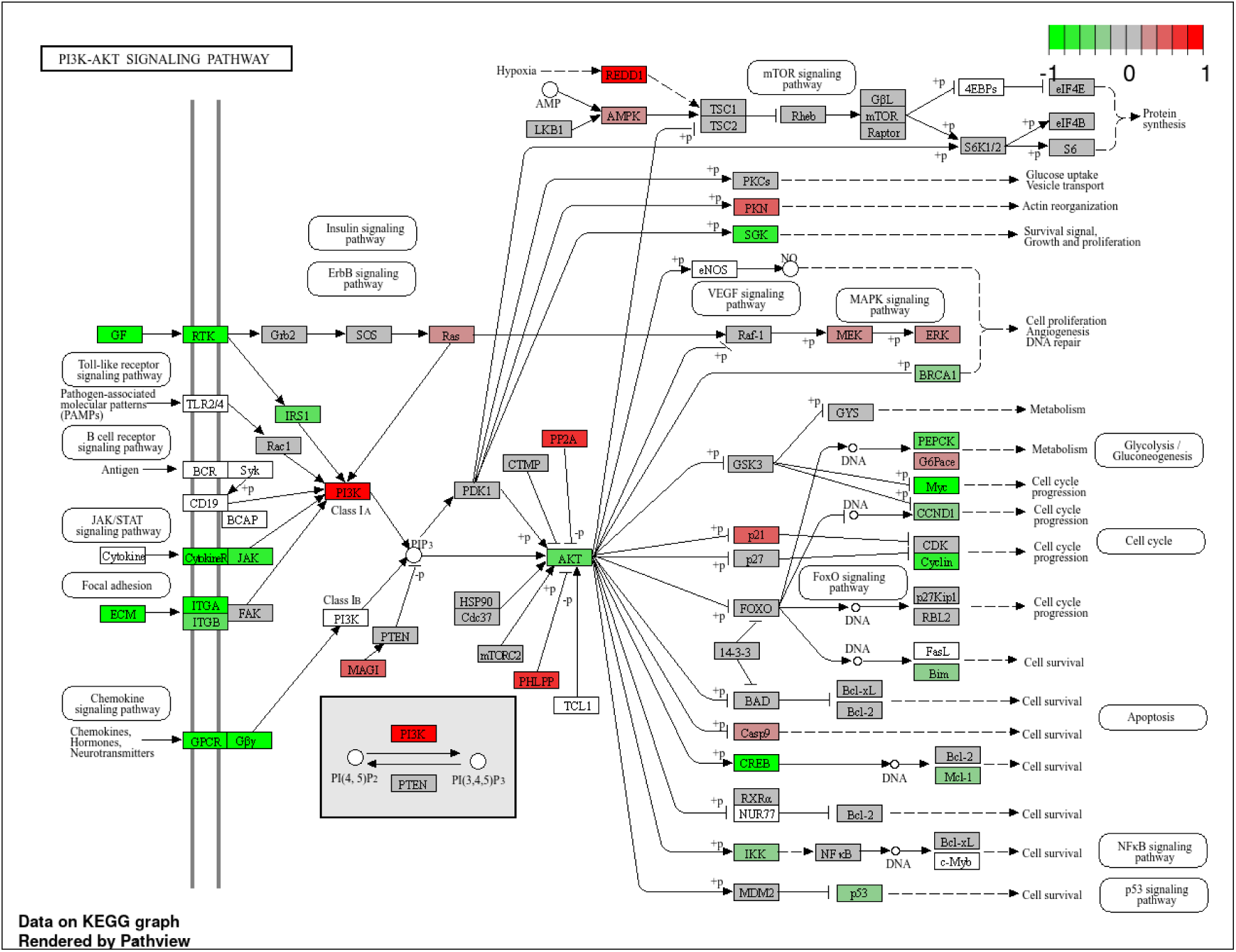
Schematic representation of the PI3K-AKT signaling pathway signaling pathways using KEGG pathways analysis for *NPHP1^ko2^* kidney organoids compared to *NPHP1^WT^*.

**Figure S13.**
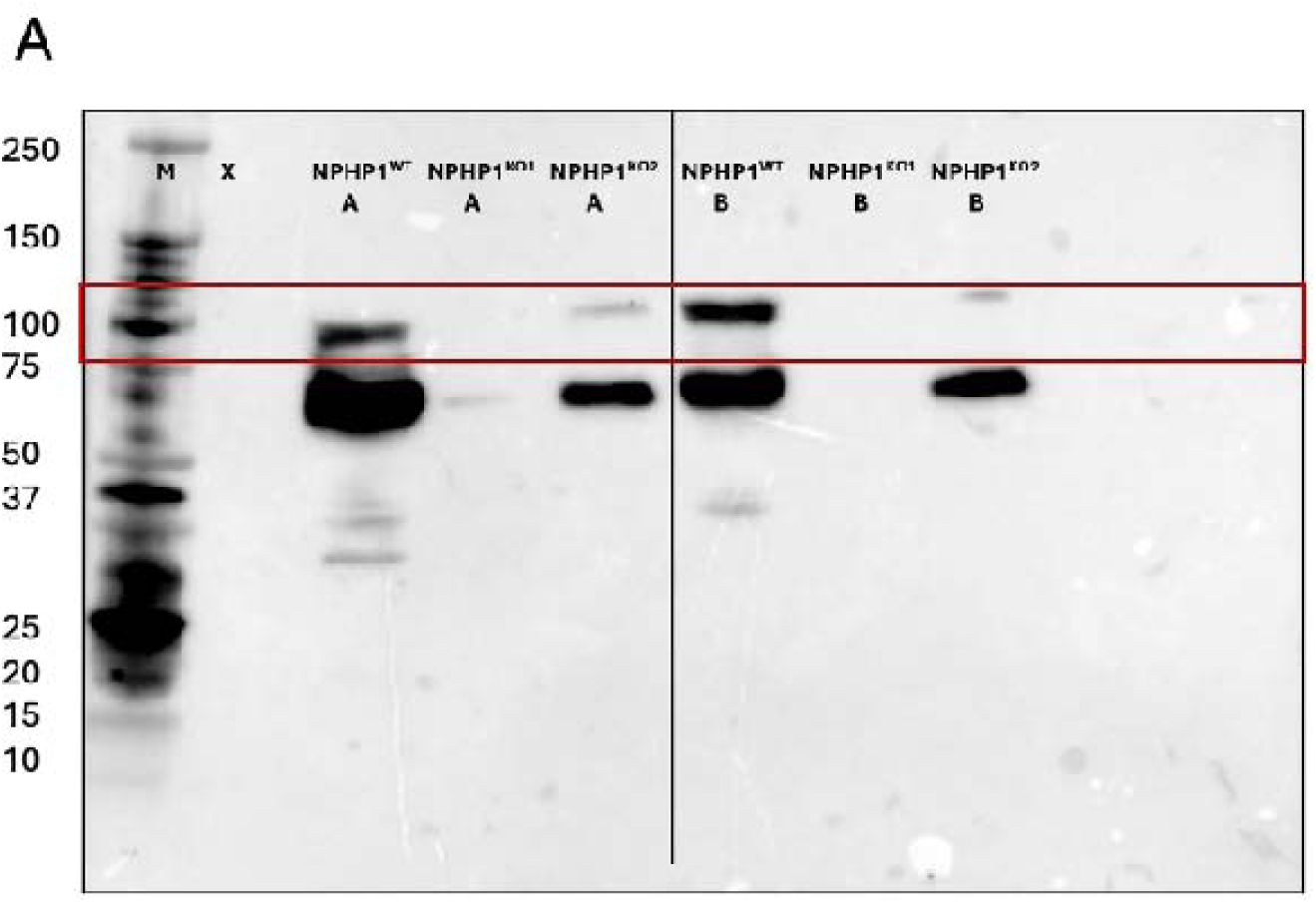
Western Blot validation of NPHP1 expression in *NPHP1*^WT^, *NPHP1*^Ko1^, and *NPHP1*^Ko2^. (A) Western blot analysis of NPHP1 expression in *NPHP1*^WT^, *NPHP1*^Ko1^, and *NPHP1*^Ko2^ samples, each generated independently from Group A and Group B differentiation protocols. The expected NPHP1 band (∼83 kDa) is indicated by the red rectangle.

**Figure S14.**
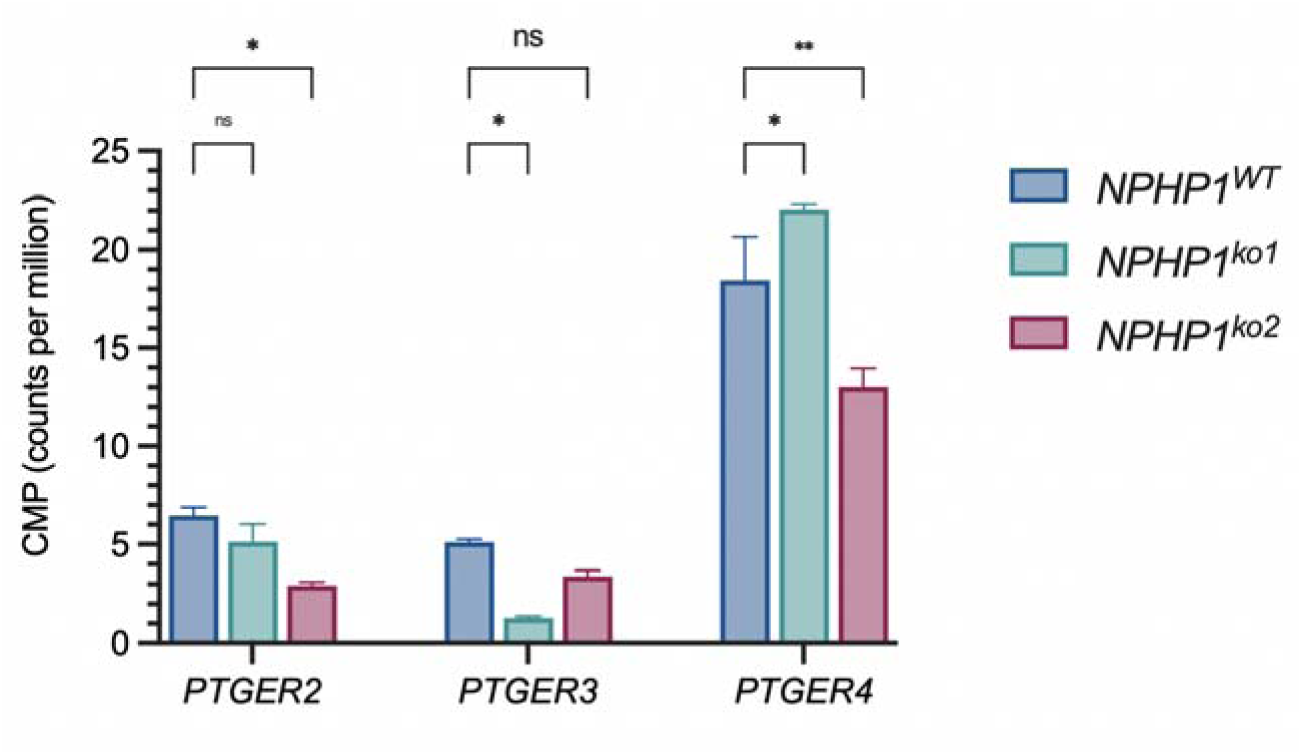
Gene expression expressed as counts per million (CPM) of prostaglandin E2 receptors in *NPHP1-* depleted kidney organoids compared to *NPHP1^WT^.* A one-way ANOVA analysis was performed (N=3, p<0.05).

**Figure S15.**
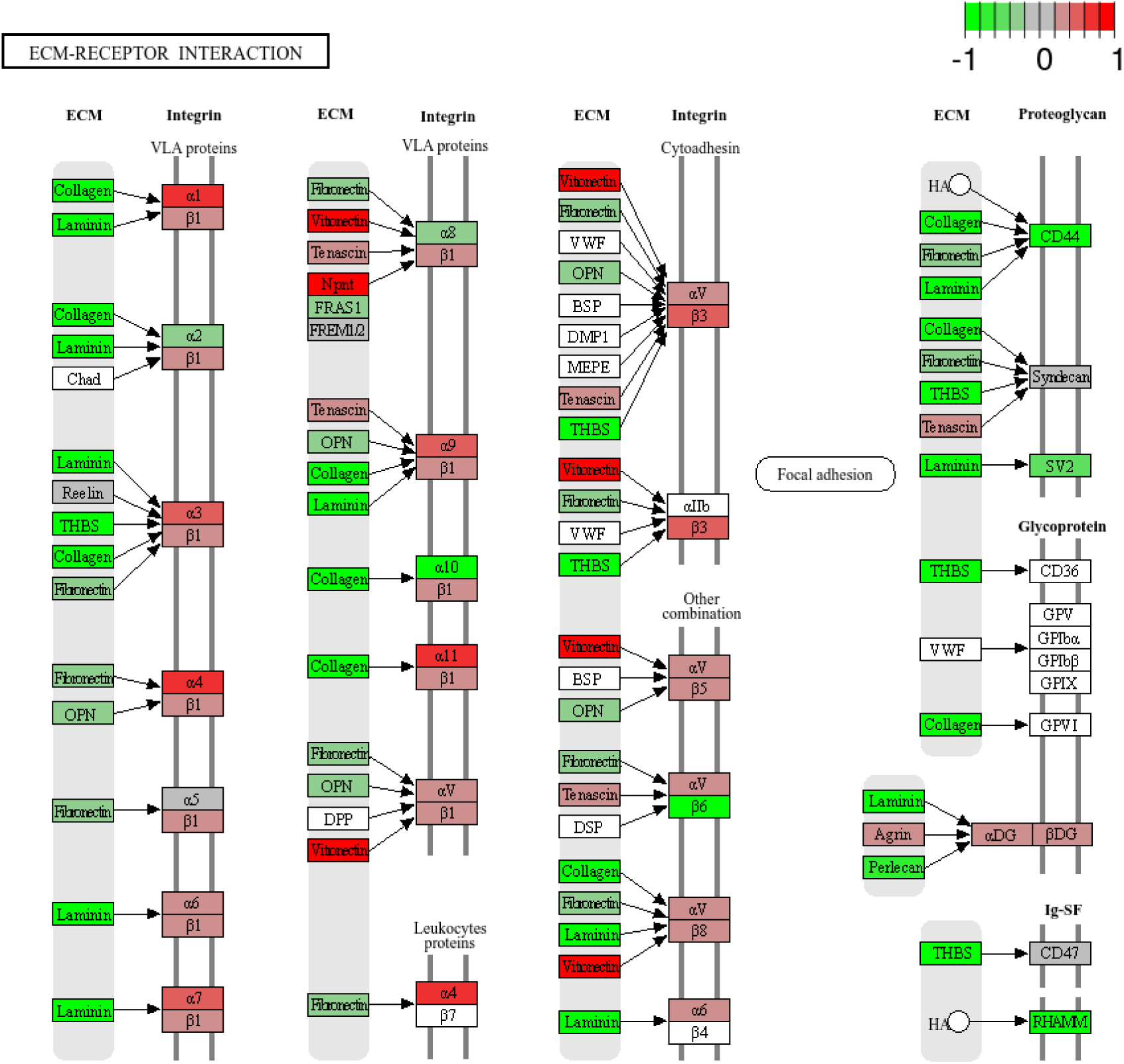
Schematic representation of the ECM-receptor interaction pathway (KEGG ID: hsa04512) using KEGG pathways analysis for *NPHP1^ko1^* kidney organoids compared to *NPHP1^WT^*.

**Figure S16.**
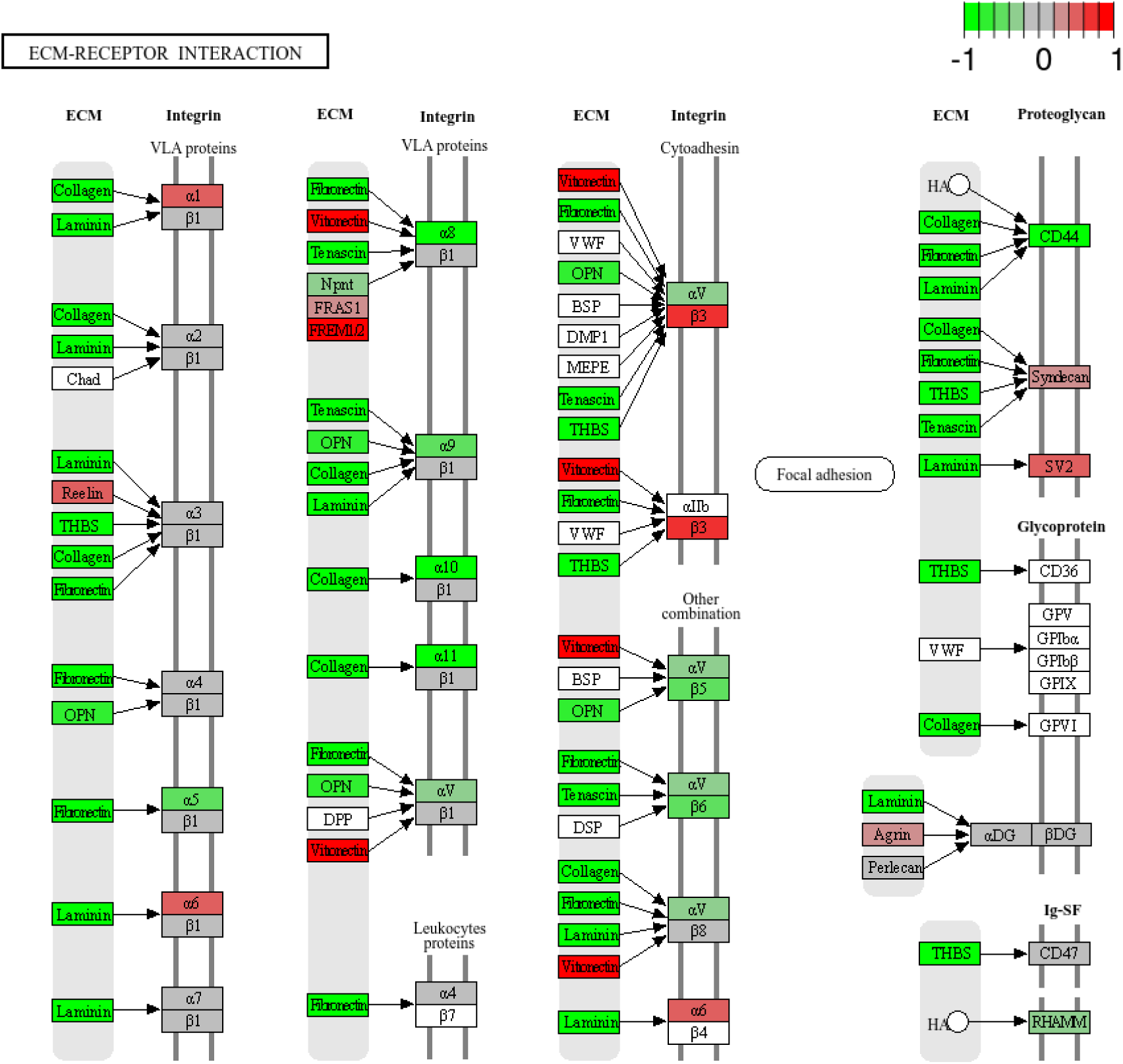
Schematic representation of the ECM-receptor interaction pathway (KEGG ID: hsa04512) using KEGG pathways analysis for *NPHP1^ko2^* kidney organoids compared to *NPHP1^WT^*.

### 8.2 Supplementary Tables (S1-S11)

**Table S1.** Nephron segment CPMs. Normalized bulk RNA-Sequencing data expressed as counts per million (CPM).

**Table S2.** Principal component analysis (PC1-5) including *NPHP1*^WT^ (WT_Untreated), *NPHP1*^ko1^ (NPHP1-C1_Untreated), and *NPHP1*^ko2^ (NPHP1-C3_Untreated) kidney organoids.

**Table S3.** Differentially expressed genes analysis (DEG1) including log2-fold changes, adjusted p-values and FDR values for *NPHP1*^WT^ (WT_Untreated), *NPHP1*^ko1^ (NPHP1-C1_Untreated), and *NPHP1*^ko2^ (NPHP1-C3_Untreated) kidney organoids.

**Table S4.** Enrichment upon differential expression analysis (DEG2) using KEGG, including top 10 up- and down-enriched pathways when comparing *NPHP1*^ko1^ to *NPHP1*^WT^ kidney organoids. The table includes the pathway names and size, number of genes enriched per pathway, the gene IDs, fold enrichments, and FDR values.

**Table S5.** Enrichment upon differential expression analysis (DEG2) using KEGG, including top 10 up- and down-enriched pathways when comparing *NPHP1*^ko2^ to *NPHP1*^WT^ kidney organoids. The table includes the pathway names and size, number of genes enriched per pathway, the gene IDs, fold enrichments, and FDR values.

**Table S6.** Principal component analysis (PC1-5) including *NPHP1*^WT^ (WT_Untreated), *NPHP1*^ko1^ (NPHP1-C1_Untreated), *NPHP1*^ko2^ (NPHP1-C3_Untreated), *NPHP1*^WT^ + UVC (WT_UVC), *NPHP1*^ko1^ + UVC (NPHP1_C1_UVC), and *NPHP1*^ko2^ + UVC (NPHP1_C3_UVC) kidney organoids.

**Table S7.** Data used to generate the Hierarchical clustering heatmap when clustering the top 1000 differentially expressed genes amongst all samples, including *NPHP1*^WT^ (WT_Untreated), *NPHP1*^ko1^ (NPHP1-C1_Untreated), *NPHP1*^ko2^ (NPHP1-C3_Untreated), *NPHP1*^WT^ + UVC (WT_UVC), *NPHP1*^ko1^ + UVC (NPHP1_C1_UVC), and *NPHP1*^ko2^ + UVC (NPHP1_C3_UVC) kidney organoids.

**Table S8.** PCA (PC1-5) including *NPHP1*^ko1^ (NPHP1-C1_Untreated) and *NPHP1*^ko2^ (NPHP1-C3_Untreated).

**Table S9.** Top1000 clustering heatmap including *NPHP1*^ko1^ (NPHP1-C1_Untreated) and *NPHP1*^ko2^ (NPHP1-C3_Untreated).

**Table S10.** Top enriched DEGs including *NPHP1*^ko1^ (NPHP1-C1_Untreated) and *NPHP1*^ko2^ (NPHP1-C3_Untreated).

**Table S11.** Top enriched pathways including *NPHP1*^ko1^ (NPHP1-C1_Untreated) and *NPHP1*^ko2^ (NPHP1-C3_Untreated).

**Table S12.** Gene expression expressed in counts per million (CPM) for the prostaglandin E2 receptors *PTGER2*, *PTGE3*, and *PTGER4*, including *NPHP1*^WT^ (WT_Untreated), *NPHP1*^ko1^ (NPHP1-C1_Untreated), *NPHP1*^ko2^ (NPHP1-C3_Untreated) kidney organoids. The gene expression of *PTGER1* is not shown since it did not pass the pre-processing filter of our RNA-Seq analysis.

